# 3D soft hydrogels induce human mesenchymal stem cells “deep” quiescence

**DOI:** 10.1101/2021.03.11.434638

**Authors:** David Boaventura Gomes, Ana Filipa Henriques Lourenço, Clarissa Tomasina, Bryan Chömpff, Hong Liu, Nicole Bouvy, Sandra Camarero-Espinosa, Lorenzo Moroni

## Abstract

Human mesenchymal stem/stromal cells (hMSCs) present a great opportunity for tissue regeneration due to their multipotent capacity. However, when cultured on 2D tissue culture polystyrene (TCPS) plates, hMSCs lose their differentiation capacity and clinical potential. It has been reported that cells need a more physiologically relevant micro-environment that allows them to maintain their phenotype. Here, we have developed a 3D alginate hydrogel functionalized with the Arg-Gly-Asp (RGD) sequence and having low mechanical stiffness that mimics the mechanical properties (>5 KPa) of bone marrow. hMSCs cultured in these hydrogels appeared to be halted in G_1_ phase of the cell cycle and to be non-proliferative, as shown by flow cytometry and 5-Ethynyl-2’-deoxyuridine (EdU) staining, respectively. Their quiescent state was characterized by an upregulation of enhancer of zeste homolog 1 (EZH1) at the gene level, forkhead box O3 (FoxO3) and cyclin-dependent kinase inhibitor 1B (p27) at the gene and protein levels compared to hMSCs grown in 2D TCPS. Comparative studies in 3D hydrogels of alginate-RGD presenting higher concentration of the peptide or in collagen hydrogels revealed that independently of the concentration of RGD or the chemistry of the adhesion motives, hMSCs cultured in 3D presented a similar phenotype.

This quiescent phenotype was exclusive of 3D cultures. In 2D, even when cells were starved of fetal bovine serum (FBS) and became also non-proliferative, the expression of these markers was not observed. We propose that this difference may be the result of mammalian target of rapamycin complex 1 (mTORC1) being downregulated in hMSCs cultured in 3D hydrogels, which induces cells to be in “deep” quiescence and be kept alive *ex vivo* for a long period of time. Our results represent a step forward towards understanding hMSCs quiescence and its molecular pathways, providing more insight for hMSCs cell therapies.

## Introduction

Adult stem cells (ASCs) are capable of providing identical daughter cells and differentiate into other cell lineages. These cells are present in almost all mammalian tissues and have unique abilities fundamental for tissue homeostasis, allowing mammalian organisms to cope with tissue lost due to normal cell turnover or tissue damage [1]. ASCs often reside in a niche, which works as a reservoir system with characteristic biophysical and biochemical properties that orchestrate the balance between proliferation and quiescence of its population. These intrinsic niche properties also determine a specific interplay between cells and extracellular matrix (ECM). Disrupting this interplay leads to ASC phenotype changes that can result in differentiation, senescence or proliferation, depleting the pool of quiescent cells [2]. In fact, most ASCs adopt a completely different phenotype when cultured *in vitro* where niche biophysical and biochemical properties are not met. For instance, hematopoietic stem cells (HSCs) have reduced proliferation and decreased viability when grown on tissue culture polystyrene (TCPS) without their native ECM [3]. Muscle stem cells (MuSC) differentiate towards muscle cells, but are not capable of keeping an undifferentiated state when cultured in TCPS [4]. Such phenotype changes have been reported for several different ASC populations in different niches. Attempts to revert their phenotype have been focused on trying to mimic the ECM properties of each stem cell niche [4]. For instance, tuning the chemistry and elasticity of the cell culture substrate have enabled HSCs to keep their stemness, and MuSCs their self-renewal phenotype. Ferreira et al. [3] have shown that covering TCPS with fibrin improved dramatically HSCs proliferation rate without compromising their primitive phenotype, while Gilbert et al. [4] have shown that MuSCs cultured *in vitro* on a soft hydrogel (12 kPa) mimicking muscle elasticity were able to self-renew.

Human mesenchymal stem/stromal cells (hMSCs), another type of ASC derived from bone marrow, have been shown to also change their phenotype upon changes to their niche properties. These cells are multipotent and show great promise in skeletal disorder therapies. Yu et al. [5] demonstrated that changes on biophysical properties of the ECM are a key modulator of the periosteal stem cell niche. When this tissue was removed from the animal’s bone and lost its *in vivo* prestress, there was a disruption of the ECM integrity, which led to an increase of round nuclei density. These changes were attributed to an epithelial to mesenchymal transition in which the cells become more proliferative and upregulate genes responsible for secretion of ECM proteins associated with tissue genesis. In fact, several studies have shown that stiffer culture substrates induce higher mitotic rates [6, 7] and tissue culture plated hMSCs will double for a limited amount of times becoming then senescent [8]. Several studies have also shown that once cultured in 3D, hMSCs become less proliferative, have a different secretome and adopt a different shape when compared to two-dimensional cultures (2D) [9–11]. This highlights the need to develop an artificial 3D niche-mimicking culture system, in order to study the *in vitro* capability of hMSCs to retain their naive phenotype as quiescent ASCs that only proliferate upon specific stimuli.

Natural and synthetic polymer hydrogels can be used to study cell behavior in an artificial 3D niche. The mechanical properties of hydrogels can be tuned to match bone marrow’s Young’s modulus. Additionally, they can also be modified to present specific biochemistries that interact with the cells [12]. One such natural polymer is alginate, a polysaccharide extracted from seaweed, which is biocompatible and bioinert, presenting only hydroxyl and carboxylic groups on its surface. The lack of functional groups that directly interact with cell surface receptors, the low protein absorption, and the ease of functionalization and cross-linking make it an ideal candidate as a naive cell culture platform [13]. Alginate also can be chemically modified to present cell-binding motifs capable of inducing desired and specific biological effects [14–16], such as enzymatically degradable peptides that promote cell proliferation or cell delivery [17].

Our aim in the present study was to design and fabricate a peptide-modified alginate 3D hydrogel mimicking key aspects of the bone marrow niche, in order to drive hMSCs towards quiescence. We mimicked the native biophysical and biochemical properties of the hMSCs’ niche using Arg-Gly-Asp (RGD) as a model peptide sequence that was introduced onto the backbone of alginate at different concentrations, resulting on hydrogels with “low” and “high” RGD content. Using these 3D systems, we induced hMSC quiescence and nonproliferation *in vitro*, as judged by flow cytometry, 5-ethynyl-2’-deoxyuridine staining, and gene and protein upregulation of quiescence-specific markers. These tools and insights may be used to better control hMSCs for therapeutic applications, and serve as a model to create artificial 3D niches for other ASC types.

## Materials and Methods

### Cell culture

Bone marrow hMSCs were isolated, with written consent from a 22-year-old male from Texas A&M Health Science Center [18] after ethical approval from the local and national authorities. Mononuclear cells were isolated by centrifugation. Isolated hMSCs were verified for their differentiation capacity and were received at passage 1. For further expansion, hMSCs were seeded at 1000 cells/cm^2^ in α-minimum essential media (αMEM) + Glutamax medium (Thermo Fisher Scientific) supplemented with 10% (v/v) fetal bovine serum (FBS) (Sigma-Aldrich) (basic medium) at 37 °C in 5% CO_2_. Upon reaching 70–80% confluence, cells were trypsinized in 0.05% Trypsin and 0.53 mM ethylenediaminetetraacetic acid (EDTA) (Thermo Fisher Scientific) and used for experiments at passage 5.

### 2D cell culture quiescence models

To establish a quiescence model, 2D cell culture systems on normal and starving conditions were studied. For that, hMSCs were seeded on TCPS at 1000 cells/cm^2^ and left to attach overnight. For the normal condition, cells were cultured in expansion medium (αMEM+Glutamax supplemented with 10% FBS). For the starving condition, cells were cultured in αMEM+Glutamax without FBS. After 3 days of culture, cells were incubated with 10 μM of EdU (modified thymidine analogue that is efficiently incorporated into newly synthesized DNA and can be fluorescently labelled), until day 6 when cells were fixed with 3.6% (v/v) paraformaldehyde (PFA) (Sigma-Aldrich) in 1× phosphate-buffered saline (PBS) for 15 min at room temperature (RT). In order to understand whether hMSCs would be able to re-enter the cell cycle, a third condition, named rescue condition, was also established where cells were left in starving condition until day 6, when the medium was switched to expansion medium (normal condition) and EdU was added. At day 8, cells were then fixed with 3.6% (v/v) PFA in PBS as previously described.

### Alginate-RGD hydrogels

Food-grade alginate kindly provided by FMC polymers, with a high guluronic acid content (74%) [14], was purified according Neves et al. [19]. In brief, alginate was dissolved in MilliQ water at a concentration of 1% w/v. Activated charcoal (Sigma-Aldrich) was added (2% w/v) and left to stir for 1 h at RT. The solution was then centrifuged at 27000 × *g* for 1 h. The supernatant was decanted and filtered through 1.2 µm, 0.45 µm and 0.2 µm pore diameter filters (Sigma-Aldrich) twice before being freeze-dried when a cotton-like solid was obtained. We grafted the RGD peptide at low and high concentrations, termed RGD-low and RGD-high respectively, to the alginate backbone. First, alginate was dissolved in 2-(N-morpholino)ethanesulfonic acid (MES) buffer (1% w/v, Sigma-Aldrich) with pH corrected to 6.5 with 0.1 M NaOH (Sigma-Aldrich). For alginate-RGD low, 48.42 mg of N-(3-Dimethylaminopropyl)-N’-ehtylcarbodiimide hydrochloride (EDC-HCl) (Sigma-Aldrich) were added to the solution along with 27.40 mg of N-hydroxy-sulfosuccinimide (sulfo-NHS) (TCI Europe) per 1 g alginate. For alginate-RGD high, 2.64 mg of EDC-HCl and 2.13 mg of Sulfo-NHS per gram were added per 1 g alginate. Each solution was stirred for approximately 30 min, then the pH was then adjusted to 8 using 0.1 M NaOH. Afterwards, 16.70 mg or 50 mg of GGGGRGDSP peptide (China peptides) per 1 g of alginate were added to produce alginate RGD-low or -high, respectively. The flask was then completely covered with aluminum foil and the solution was left to stir overnight (approximately 18 h) at RT when it was quenched with 18 mg hydroxylamine hydrochloride (Sigma-Aldrich). The solution was dialyzed with molecular weight cut-off membranes (SnakeSkin™ Dialysis Tubing, ThermoFisher-Scientific, 10 kDa MWCO, 22 mm) against a decreasing NaCl (Sigma-Aldrich) gradient, with a starting concentration of 6 g/L, for 3 days. Afterwards, the solution was freeze-dried for another 3 days and a cotton-like solid was obtained.

### NMR analysis

Samples of alginate functionalized with RGD were prepared by dissolving 3–4 mg of the product in 750 μL of sodium acetate-d_3_ solution in H_2_O (pH 3). NMR spectra were recorded on a 700 MHz Bruker spectrometer operating at 325K and applying a water suppression pulse sequence.

### Cell encapsulation in hydrogels

After hMSCs trypsinization from the flasks, a pellet was obtained by centrifuging the cell suspension at 300 × *g* for 5 min in a 1.5 mL conical tube. After discarding the supernatant, 1% w/v alginate in 0.9% (g/v) NaCl solution, sterilized through a 0.22 μm pore filter (Sigma), was added to the tube and mixed with the cells, at 10 × 10^6^ cells per mL. The cells and the hydrogel were mixed through resuspension with a gel pipette (Gibco), to avoid shear stress. The hydrogels were left in the solution for 5 min and replaced by 1 mL of cell culture medium for 30 min, until it was replaced again for the experiment. Collagen hydrogels were prepared at 6 mg/mL. For a total of 300 μL, the following reagents were added to a 1.5 mL microcentrifuge tube in the following order: 30 μL of PBS 10×, 4.96 uL of 1 M NaOH (VWR), 49.21 μL of ultrapure water, and 215.8 μL of rat tail high concentration collagen type I at 8.34 mg/mL (Corning). After mixing well, the final solution was added on top of a pellet of 3 × 10^6^ hMSCs to achieve a final cell density of 10 × 10^6^ cells/mL. Individual 10 μL collagen solutions were pipetted on top of a non-adherent, 24-well plate and allowed to gel at 37 °C for 30 min with water in the plate to provide humidity to the system. After full gelation, the hydrogels were scooped with a spatula and dropped into 1 mL of cell culture medium. The medium was changed every other day throughout the time in culture. After the experiments were completed, the hydrogels were washed twice with 1× Tris-buffered saline (TBS) and fixed with 3.6% (v/v) PFA in TBS for 15 min at RT. The samples were then kept at 4 °C until further use.

### Rheological measurements

For alginate samples, a cast was made consisting of a 1 mm thick slab of PET with 8 mm diameter holes, which was placed on top of a glass slide covered in write-on tape, to ensure partial absorption and a homogeneous spread of the crosslinking solution. A crosslinking solution (1 mL of 0.1 M CaCl_2_) was pipetted in between the slide and mold, allowing it to spread under the mold. The alginate solutions (51 µL) were then pipetted into the holes until the edges of the drop came into contact with the layer of crosslinking solution. The alginate was then left to crosslink for 3 h. CaCl_2_ crosslinking solution was pipetted onto the setup every hour to counteract evaporation. Flat discs were obtained with 0.5 mm thickness on 24-well plates and subsequently punched to 8 mm diameter.

Rheological measurements were performed on a TA Instruments at 25 °C using a parallel flat plate geometry with top and bottom diameters of 8 mm and 50 mm, respectively. Samples were loaded from the cast onto the loading dock. The top plate was then lowered slowly onto the gel until a pre-load normal force of 0.5 N was registered, ensuring a complete fill up of the space between the plates. The loading dock was then covered in TBS and the gel was allowed to relax back to a normal force of 0.1 N. To determine the linear viscoelastic region (LVR), an oscillatory amplitude sweep was conducted at 1 Hz frequency and from an amplitude of 0.1% strain to a maximum of 100%. Five points per decade were recorded during the experiment. The storage (G’) and loss (G’’) moduli were graphed as a function of the oscillation strain. A constant strain for the frequency sweeps was then selected within the LVR. This process was conducted twice for verification. Oscillatory frequency sweeps were performed at a constant strain, determined within the LVR for each condition and each batch, with a frequency range between 0.1 and 100 Hz. Five points per decade were recorded during performance. G’ and G’’ were then graphed as a function of strain and frequency, respectively.

### hMSCs lineages differentiation

After cell culture in the different 2D TCPS conditions, hMSCs were trypsinized and centrifuged at 500 × *g* for 5 min to be seeded for lineage differentiations. For alginate-RGD low 3D culture, each hydrogel was placed on a propylene 15 mL tube and washed with 5 mL PBS. After discarding the PBS, alginate-RGD low hydrogels were incubated with 5 mL of 50 mM EDTA for 5 min, until the hydrogels dissolved. The suspension obtained was then aspirated up and down through a 30 G needle to ensure total hydrogel disruption. The cell suspension was then centrifuged at 800 × *g* for 10 min and resuspended in medium to be seeded.

For osteogenesis, cells were plated at 5000 cells/cm^2^ in a Falcon polystyrene 24-well microplate (Fisher Scientific) in αMEM + Glutamax with 10% FBS and 1% penicillin-streptomycin (P/S) (Thermo Fisher) for 7 d until reaching full confluency. Cells were then cultured in osteogenic medium, with every other day medium changes supplemented with 200 μM L-Ascorbate2-phosphate (Sigma-Aldrich) and 10 nM dexamethasone (Sigma-Aldrich) for 21 d. After 21 d, β-Glycerol phosphate (BGP) was added to the medium to a final concentration of 10 mM to accelerate mineralization for an additional 7 d. Cells were then fixed with 3.6% (v/v) PFA in PBS for 15 min at RT and stained with alizarin red for imaging. Briefly, well plates were washed two times with MilliQ H_2_O and then stained with 400 μL of 2% w/V alizarin red S (VWR) for 10 min. After being washed two times with MilliQ H_2_O for 5 min at 50 rpm in an orbital shaker, the samples were imaged.

For adipogensis, cells were plated at 5000 cells/cm^2^ in a Falcon™ polystyrene 24-well microplate in αMEM + Glutamax supplemented with 10% FBS, 1% P/S, 0.2 mM indomethacin (Sigma-Aldrich), 1 μM dexamethasone, 0.5 mM 3-Isobutyl-1-methylxanthine (Sigma-Aldrich), and 2.5 μg/ml insulin (Sigma-Aldrich). Medium was replaced every other day. Cells were then fixed with 3.6% (v/v) PFA in PBS for 15 min at RT and stained with oil red O. Briefly, the plates were washed with ddH_2_O, incubated with 60% isopropanol (v/v) (VWR) in water for 5 min, and then in oil red O (Sigma-Aldrich) working solution (3 parts oil red O at 3 mg/mL in isopropanol with 2 parts ddH_2_O) for 20 min. Then, the plates were carefully washed two times with ddH_2_O until imaging was completed.

For chondrogenesis, 250,000 cells were pelleted with αMEM + Glutamax at 450*g* for 8 min at RT in 15 ml polypropylene conical tubes and incubated for 24 h with the lid partially opened to allow gas exchange. After pellet formation, cells were cultured in the same tubes in 0.5 mL of high glucose DMEM (Thermo Fisher) supplemented with 100 µg/ml sodium pyruvate (Thermo Fisher), 100 U/mL P/S, 100 μg/ml insulin-transferrin-selenium (ITS) premix (Sigma-Aldrich), 0.2 mM ascorbate-2-phosphate, 100 nM dexamethasone, 40 μg/ml L-proline (Sigma-Aldrich) and 0.01 μg/ml transforming growth factor-β3 (TGF-β3) (Peprotech). The medium was replaced every other day. After 28 d of culture, cells were fixed with 3.6% (v/v) PFA for 15 min at RT in PBS and pellet sections stained with Alcian blue (Sigma-Aldrich). Briefly, pellets were processed for paraffin embedding (described more in detail in the *in vivo* subsection) and the blocks were cut into 3 µm slices. The samples were then deparaffinized and hydrated to ddH_2_O. Samples were incubated in 1% Alcian blue 8 GX (Sigma-Aldrich) in 3% (v/v) acetic acid (Sigma-Aldrich) solution for 30 min. After being washed twice with running tap water, the samples were then counterstained with 0.1% (w/v) nuclear fast red in water solution for 5 min. The samples were rehydrated, cleared with Neo-Clear solution (xylene substitute) (Merck), mounted with DPX mounting medium (Sigma-Aldrich), and stored until imaged.

### Cell viability and metabolic activity

Cell viability was assessed with LIVE/DEAD™ Viability/Cytotoxicity Kit (Thermo Fisher) following the manufacturer instructions. Briefly, cells were washed with PBS (TCPS samples) or TBS (hydrogel samples) twice. A mixture of 6 µM ethidium homodimer and 1 µM calcein acetoxymethyl (calcein AM) was added to the wells and incubated for 30 min protected from light. The samples were then washed with PBS (TCPS samples) or TBS (for hydrogels) and left with medium until imaged under the microscope.

For cell metabolic activity, PrestoBlue™ cell viability reagent (Thermo Fisher) was used. Briefly, cells were washed with PBS (TCPS samples) or TBS (hydrogel samples) twice. Then, 500 µL of a 1:10 diluted PrestoBlue solution was added to the wells and incubated for 2 h protected from light; 100 µL of the incubated solution were then pipetted in each of four wells of a black 96-well plate. In a CLARIOstar® Plus plate reader (Isogen), the samples were excited at 560 nm and fluorescence was measured at 590 nm.

### EdU staining

For EdU staining, a Click-iT™ EdU Cell Proliferation Kit for Imaging, Alexa Fluor™ 647 dye (Thermo Fisher) was used. Briefly, cells were fixed and washed twice with 1× PBS, and permeabilized with 0.5% (v/v) Triton X-100 (Merck) in PBS for 20 min at RT. After being washed twice with PBS, cells were then incubated in the Click-iT® reaction cocktail (prepared according to the manufacturer manual) for 30 min at RT, protected from light. The cocktail was then removed, cells washed twice with PBS and counterstained with 10 μg/mL Hoechst33342 in 1× PBS for 10 min at RT, protected from light. Images were then taken on a wide-field fluorescence microscope (Nikon-Ti) and analyzed with Fiji software.

### Cell culture under hypoxia conditions

Immortalized MSCs (iMSCs) cultured in 2D TCPS in hypoxia, 1% (v/v) O_2_ incubator (used as control condition), and hMSCs encapsulated in 3D alginate-RGD low hydrogels in normoxia, 20% (v/v) O_2_ incubator, were cultured for 7 d. To evaluate the hypoxic conditions, the Image-iT™ Hypoxia Reagent kit (Thermo Fisher) was used according to the manufacturer instructions. In brief, samples were incubated with 50 μM of Image-iT™ Hypoxia Reagent stock solution for 1 h at 37 °C, washed with warm medium and incubated for 2 more hours with medium at 37°C. The samples were imaged immediately after, with excitation at 490 nm and emission at 610 nm on a SP8 Leica confocal microscope.

To assess proliferation under hypoxic conditions, hMSCs were cultured for 7 d in 2D TCPS either in hypoxia or normoxia. At 24 h prior to fixation with 2.6% PFA in PBS, EdU was added to the medium. Proliferation was then assessed after EdU staining, as described above.

### Immunofluorescence and imaging

After fixation, cells were permeabilized as explained above. Then, the samples were blocked with a solution of 1% bovine serum albumin (BSA) and 4% goat serum (GS) in PBS (for samples without hydrogels) or TBS (for samples with hydrogels) for 1 h at RT. Primary antibody Anti-Ki67 (Fisher Scientific, ref no. 11357433) at 1:200 dilution, anti-nestin (ThermoFisher, ref no. MA1-110) at 1:100 dilution, anti-stro-1 (ThermoFisher, 39-8401) at 1:100 dilution, recombinant anti-p27 KIP 1 antibody [Y236] (Abcam, ab32034) at 1:100 dilution, and RB1 (Abcam, ab181616) at 1:500 dilution, anti-YAP1 (Abcam, ab52771) at 1:250 dilution, anti-paxillin (Abcam, ab32084) at 1:1000 dilution, or anti-zyxin (Abcam, ab58210) at 1:1000 dilution were incubated overnight at 4 °C. The samples were then washed three times with PBS and incubated with anti-mouse or anti-rabbit Alexa Fluor 488 or 568 (Thermo Fisher Scientific) for 1 h at RT protected from light, and overnight for anti-zyxin and anti-paxillin staining. Images were taken on a wide-field fluorescence microscope (Nikon-Ti) or on an SP8 confocal (Leica) microscope. All images within an experiment were captured on the same day and using the same settings, to allow for quantitative comparisons.

For YAP1 quantification, the nucleus and an area in the cytoplasm were selected and the overall pixel intensity in each area was measured using Fiji. The ratio between nuclear and cytoplasmic intensity was then calculated per cell. For zyxin and paxillin, the number of zyxin and paxillin dots inside of the cytoplasm were counted and divided by the total cell area.

### RNA isolation and qPCR

hMSCs were cultured in 2D TCPS conditions or in alginate-RGD low hydrogel for 6 days. TRIzol™ (Fisher Scientific, 1 mL) was added to each condition (175 cm^2^ TCPS flasks and four 10 μL hydrogels), and cells were dissolved and detached with the help of a cell scraper (VWR). The samples were then frozen at −80 °C until further use. For RNA isolation, samples were thawed on ice and 200 μL of chloroform (Sigma-Aldrich) were added. After shaking vigorously for 15 s, the samples were left for 15 min at RT and then centrifuged at 12000 × *g* for 15 min at 4 °C. The upper aqueous layer was recovered and 500 μL of isopropanol (VWR) were then added. The samples were incubated for 10 min at RT and centrifuged at 12000 × *g* for 10 min at 4°C. All but 200 μL of supernatant was discarded and a same volume of molecular-grade ethanol (Thermo Fisher Scientific) was mixed into the solution with a pipette to make a lysate. The samples followed then the protocol established by RNeasy Mini Kit (Qiagen) for total RNA isolation. The RNA concentration was quantified with a nanodrop and the samples concentration were evened with nucleoside-free water (Bio-Rad). To synthesize cDNA from RNA, iScript™ cDNA Synthesis Kit (Bio-Rad) was used following the manufacturer’s instructions with a reaction of 5 min at 25°C, 20 min at 46°C, 1 min at 95°C. Samples were then frozen at −30°C for storage or in ice for further use.

### Quantitative real-time RT-PCR

Primers for *Cyclin A_2_*, *Cyclin B_1_*, *Cyclin E_1_*, *FoxO_3_*, *p27*, *EZH1,* and *RPS27A* were selected and confirmed for their affinity to the gene with BLAST (Supplementary Table 1). For reverse transcription, 1 μg of sample was loaded with 0.3 μL of primers solution and 7.5 μL of iQ SYBR Green Supermix (Bio-Rad) diluted with RNA-free water to a total of 15 μL. After loading each reaction in a 96-well plate, the reaction was carried on a real-time PCR detection system (Bio-Rad) and following a protocol of 3 min at 95 °C and 15 sec at 95 °C, 30 sec at 55 °C (repeated 39 times), and melting curve from 65 °C to 95 °C at 0.5 °C increments for 5 min. The relative level of mRNA expression was calculated using the ΔΔCt method with *RPS27A* as a reference gene.

### Cell cycle analysis

Alginate hydrogels were washed with PBS and incubated with 50 mM EDTA in PBS for 10 min at 37 °C. After dissociating the alginate hydrogels, the cells were washed twice with ice-cold PBS. The cells were centrifuged at 500 × *g* and the PBS was aspirated. Subsequently, ice-cold absolute ethanol was added dropwise to the cells while vortexing. Cells were fixed overnight at 4 °C. Fixed samples were washed twice with PBS and resuspended in PBS. The samples were then treated with 10 μg/mL RNase A (Invitrogen) and stained with 40 μg/mL propidium iodide (Sigma-Aldrich) overnight at 4 °C in the dark. DNA content was determined by flow cytometry (BD Accuri C6). A minimum of 10,000 events was acquired per sample. The percentage of cells in different phases of the cell cycle was assessed using the FlowJo software.

### Caspase 3/7 activity assay

Caspase 3/7 activity was measured using the Caspase-Glo 3/7 assay (Promega). Caspase 3/7 assay solution was mixed 1:1 with αMEM without phenol red (ThermoFisher Scientific) (caspase 3/7 lysis buffer) and added to the cells at the moment of harvest. After 30 min of incubation, light intensity was measured at 520 nm on a CLARIOstar® Plus plate reader.

### DNA quantification

hMSCs were washed 2× with PBS to remove dead cells and medium then stored dry at −80 °C until further use. Samples were freeze-thawed twice before addition of either RLT lysis buffer (Qiagen) or the caspase lysis 3/7 buffer. Samples were freeze-thawed three times again in lysis buffer (after caspase 3/7 assay, if applicable) to ensure full lysis. TCPS samples were scraped and hydrogels were left in the lysis buffer. Samples were then diluted 50× in Tris-EDTA buffer (10 mM Tris-HCl, 1 mM EDTA, pH 7.5 (Sigma-Aldrich)) and a DNA standard curve was made in the same final solution (2% RLT or caspase lysis 3/7 buffer in Tris-EDTA buffer). A Pico green assay (ThermoFisher Scientific) was used to quantify DNA, according to the manufacturer’s protocol.

### Protein isolation and Western blot

Proteins were isolated from cells cultured on 2D TCPS or 3D hydrogels. For 2D TCPS samples, after washing the plates with cold PBS twice, 100 μL of radio-immunoprecipitation assay (RIPA) buffer (Sigma-Aldrich), supplemented with cOmplete™, Mini, EDTA-free Protease Inhibitor Cocktail (Sigma-Aldrich) were pipetted onto the plate surface and scraped with cell scrapers to ensure cell lysis. For alginate hydrogels, the same protocol described above to retrieve cells from alginate-RGD low was used. To get sufficient proteins, 24 hydrogels were used to get a single cell pellet. Experiments were repeated 3 or 4 times for replicates. Subsequently, hMSCs were resuspended in 5 mL of PBS and centrifuged at 2500 × *g* for 4 min. Supernatant was discarded and the pellet was then resuspended in 300 μL of RIPA buffer and incubated for 1 h on ice. The obtained solution was then sonicated (Qsonica, LLC) on ice for 30 s at 50% power. The samples were centrifuged for 1 min at 14,000 × *g* at 4 °C. For collagen, to get sufficient proteins, 24 hydrogels were used to get a single cell pellet by incubating the hydrogels with collagenase type IV (Stem Cells Technologies) at 1 mg/mL for 10 min. The final cell suspension was then centrifuged at 500 × *g* for 5 min. The cells were then resuspended in 5 mL of PBS and centrifuged at 2500 × *g* for 4 min. Supernatant was discarded and the pellet was then resuspended in 300 μL of RIPA buffer, incubated for 1 h on ice, and followed the same protocol for sonication as for the alginate samples. All samples were then stored at −80°C until protein quantification.

Protein quantification was done using the Pierce BCA protein assay kit (Thermo Fisher Scientific). Fourteen micrograms of protein were incubated with Laemmli loading buffer (Bio-Rad) and 10% 2-mercaptoethanol (Sigma-Aldrich) for 5 min at 95 °C, loaded into a 4–15% polyacrylamide gel (Bio-Rad), and left to run for 1 h at 100V. Proteins were transferred to a 0.45 μm polyvinylidene fluoride (PVDF) membrane (Bio-Rad) using the wet transfer method for 1 h at 100V. Membranes were blocked for 1 h with 5% nonfat milk powder (Bio-Rad) in TBS + 0.05% Tween-20 (Sigma-Aldrich) for anti-RB1 (Abcam, ab181616, 1:2000), anti-mTOR (Abcam, ab134903, 1:2000), recombinant anti-p27 KIP 1 antibody [Y236] (anti-p27) (Abcam, ab32034, 1:1000), anti pan-akt (phospho T308) (anti-Akt) (Abcam, ab38449, 1:500), anti-GAPDH (Santa Cruz, sc-365062, 1:1000), or anti-RICTOR (Abcam, ab219950, 1:500), and with 2% BSA in PBS + 0.05% Tween-20 for anti-FoxO3 (Abcam, ab109629, 1:1000) and anti-RAPTOR (Abcam, ab40768, 1:5000). Anti-p27, anti-GAPDH, anti-Akt and anti-FoxO3 primary antibody incubations were performed for 48 h, while the rest were performed overnight at 4 °C in the respective blocking buffer. Blots were subsequently incubated with 1:2000 dilution of goat-anti-rabbit or mouse horseradish peroxidase (Bio-Rad) in blocking buffer for 1 h at RT. Protein bands were then visualized using Clarity Western ECL (Bio-Rad). Quantifications of band intensity were done in Fiji using the gel quantification tool.

### Actin-myosin and ROCK inhibition

To test the effect of actin-myosin tension, Rock and MRTF on hMSCs quiescence, cells were cultured in TCPS (2D) for 5 days in normal medium (αMEM+10% FBS). Then 100 µM blebbistatin (Sigma-Aldrich), 10 µM Y27632 (Sigma-Aldrich) (Rock inhibitor), and 12.2 µM CCG-203971 (MRFT inhibitor) (Sigma-Aldrich) were added to the medium and incubated for 24 h. Protein lysates were they isolated by following the protocol described above.

### In vivo subcutaneous implantation

All experiments and protocols were approved by the local national authority, central committee for animal experiments (in Dutch: centrale commissie dierproeven). Rats were housed at 21 °C with a 12 h light/dark cycles and had ad libitum access to water and food. Before the operation (at least 30 min prior to anesthesia), buprenorphine (0.05 mg/kg bodyweight) and carprofen (4 mg/kg) were administered to 8 female rats (Crl:NIH-Foxn1rnu, 8–10 weeks old, 140–212 g) (Charles River). Animals were anesthetized under 3–4% (v/v) isoflurane, mixed with oxygen and air; anesthesia was maintained by 2% (v/v) isoflurane, adjusted according to the clinical signs during surgery. After shaving and sterilization of the animal, four 1 cm– long linear incisions parallel to the spine were made (two in each side). Four pockets of maximum 10 mm × 10 mm were created with the help of scissors, after which 100 μL alginate-RGD low hydrogel disks, 12 mm diameter, with 10 million hMSCs per mL cultured for 24 h, were implanted inside the pockets. After implantation, the skin was closed intracutaneously by using an absorbable suture (Monocryl 4-0) and analgesics (0.03 mg/kg Buprenorphine) were administered 8 h after the operation. The morning of postoperative day 1 and 2, each animal was administered with a dose of 4 mg/kg carprofen. Thereafter, all animals were evaluated on a daily basis with a discomfort logbook score system for 21 days. At 48 h before euthanasia, 40 mg/kg EdU was administered through an intraperitoneal injection. Animals were sacrificed with gradual CO_2_ overdoses. The pocket with the sample and surrounding tissues were collected and processed for histology.

### Tissue preparation and in vivo proliferation quantification

Tissue explants were cut with surgical scissors to fit the dimensions of the silicon molds (2 × 2 × 2 cm), without disrupting the generated pocket that held the implants. Samples were then placed in 50 mL centrifugation tubes and fixed for 24 h at 4 °C in a 3.6% (v/v) solution of PFA in TBS. After fixation, the samples were transferred to new tubes and underwent embedding for 24 h each in a series of: 30% (w/v) sucrose, then 50:50 volume ratio of 30% sucrose and optimal cutting temperature (OCT, Scigen) compound, and lastly OCT only. The samples were maintained in OCT until freezing. Tissue explants were placed inside silicon molds and the molds were filled with OCT. Freezing was conducted on the liquid-vapor interface of a liquid nitrogen tank to avoid formation of bubbles. Sections of 7 µm thick were cut with a cryotome and samples were stained with hematoxylin and eosin (H&E). Sections were hydrated in de-ionized water and placed in Gill’s hematoxylin (III) (Sigma-Aldrich) for 5 min, in running water for 5 min, dehydrated and counterstained with alcoholic eosin Y (Sigma-Aldrich) for 1 min. Sections were dehydrated in 100% ethanol, allowed to air dry, and mounted in DPX.

To quantify proliferation, samples were stained for EdU. Briefly, slides were taken from −30 °C and left to stabilize at RT for 5 min. Then, rehydration was achieved by incubating them in TBS for 5 min. Following the same protocol explained above, the samples were permeabilized, incubated with EdU staining solution, and counterstained with Hoechst (1:1000) for 10 min or with Gill’s hematoxylin following the same protocol above. Tissue sections were then mounted with Fluoroshield™ (Sigma-Aldrich) and imaged on a wide-field fluorescence microscope (Nikon-Ti). The percentage of EdU-positive cells was quantified by counting all EdU-positive cells and dividing by the total number of cells inside the hydrogel area.

### Statistics

The number of biological replicates and statistical tests used are stated in the figure captions. At least three biological replicates were used for each assay. Cells selected for quantification of cell size and actin stress fibers were selected randomly. Normal distribution of each data set was tested using the Shapiro-Wilk test. For multiple comparisons within one experiment, a One-way ANOVA with Tukey’s post hoc was performed, or Kruskal-Wallis with Dunn’s post hoc as nonparametric equivalent. Experiments with one comparison were tested using a two-tailed student’s t-test. Significance was set at p<0.05. Statistical analysis was performed using GraphPad Prism 8.

## Results

### Starvation leads to hMSCs quiescence

ASCs are characterized by the ability to differentiate into different cell types, being able to be quiescent and ready to re-enter cell cycle when needed. Despite being found throughout the human lifespan, hMSCs are not considered to be typical ASC, as there is no evidence that they are able to become quiescent and re-enter the cell cycle. It was previously shown that fibroblasts are cell cycle–arrested when deprived of FBS [20], which led us to hypothesize that FBS deprivation could also induce hMSCs’ cell cycle arrest. To test this, we cultured hMSCs in normal medium with FBS or starving medium without FBS, then monitored cell proliferation by EdU staining (Figure 1A). After day 6 of culture, EdU-positive nuclei were counted and normalized to the total nuclei (Figure 1C). Almost no EdU+ cells (0.3 ± 0.3%) could be found in the starving condition (Figure 1B, C) compared to 21.9 ± 5.7% of EdU+ hMSCs grown in normal medium (Figure 1C). Immunofluorescence staining of Ki67, a marker for proliferation (Figure 1D), confirmed these observations: normal culture conditions showed ki67 colocalizing on the nuclei whereas starving conditions showed no stained cells.

**Figure 1:**
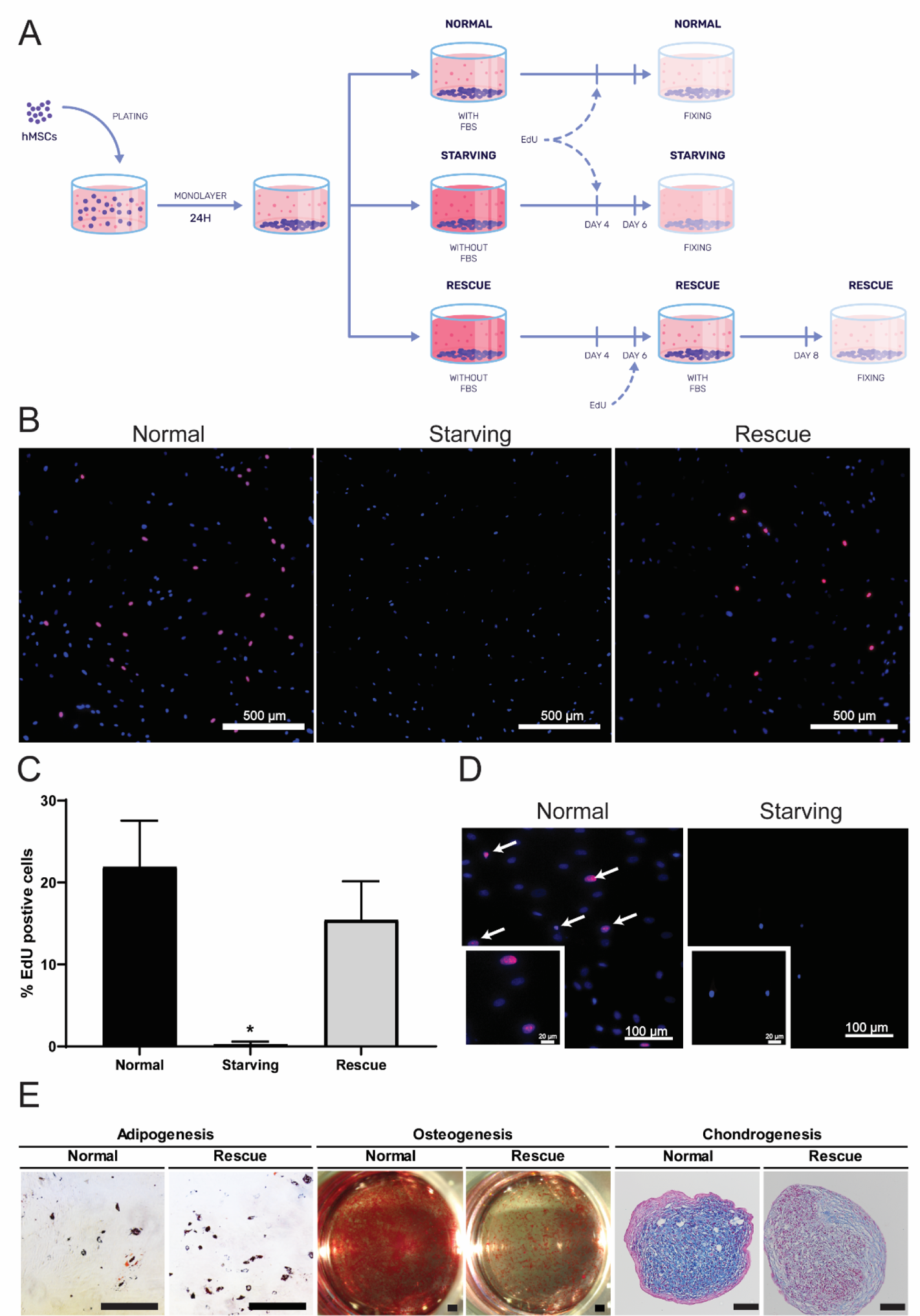
Starvation induces hMSC quiescence. (A) After 24 hours of cell adherence, hMSCs were treated with normal medium (with 10% FBS) and with starving medium (no FBS). After 3 days of treatment, cells were incubated with EdU for 2 days, until day 6 when they were fixed. For Rescue condition, cells were starved (i.e. left in Starving condition) until day 6 and medium was switched to expansion medium (i.e. left in Normal condition) with EdU. At day 8, cells were then fixed. (B) EdU immunofluorescence images of hMSCs cultures in Normal, Starving and Rescue conditions after 24 hours EdU incubation. In blue nuclei (DAPI), and in red proliferative cells (EdU), n=3; scale bar 500 μm. (C) %EdU positive cells grown for 7 days on top of TCPS Normal, Starving and Rescue 2D conditions, n=3. One-way ANOVA with multiple comparisons (*p<0.05). (D) ki67 immunofluorescence staining on hMSCs cultures on TCPS Normal and starving 2D conditions for 7 days. Nuclei stained in blue (DAPI) and proliferative cells in red (ki67), n=3. (E) After being culture for one week in Normal and Rescue conditions in 2D, hMSCs were replated and expanded in 2D TCPS and underwent adipogenic, osteogenic and chondrogenic differentiations for 21, 28 and 28 days, respectively. Bright field images show staining for oil red O, alizarin red and alcian blue for adipogenic, osteogenic and chondrogenic differentiations, respectively. n=3; Scale bars: 100 μm.

To determine whether quiescent hMSCs could re-enter the cell cycle, we performed a rescue experiment, maintaining cells in starving medium until day 6, then adding normal medium supplemented with EdU for 48 h (Figure 1A) and counted EdU-positive nuclei. The re-introduction of FBS allowed hMSCs to re-enter in the cell cycle, as seen by an increase in EdU-positive cells (15.4 ± 4.7%; Figure 1B–C) compared to starved, non-proliferative hMSCs at day 6.

To assess whether hMSCs retain their multipotency after being quiescent, cells were starved and rescued, and further expanded. Cells retained their potential to differentiate into adipocytes, osteoblasts, and chondrocytes, as demonstrated by oil red O, alizarin red, and alcian blue staining, respectively (Figure 1E). However, the rescued cells showed less efficient differentiation to osteoblasts and chondrocytes compared to normally cultured hMSCs. For example, the red reticular staining characteristic of hMSC differentiation towards osteoblasts was less intense in the rescue condition compared to the strong Alizarin staining in the normal condition. For chondrogenesis, the pellet was mostly stained in blue, characteristic of glycosaminoglycans (GAGs), for the normal condition whereas only some GAGs staining and an intense red color characteristic of fibrotic cartilage were observed in the rescued hMSCs.

Cells that were in normal and rescue conditions were also cultured under basal media (αMEM with 10% FBS without growth factors for differentiation) to assess whether the FBS-deprived quiescence could influence hMSCs to differentiate towards any specific linage. Histological staining showed that cells remained undifferentiated when no chemical input was added (Supplementary Figure 1A–C).

### Culture in 3D alginate-RGD low hydrogels induces hMSCs quiescence

Next, we tested whether hMSC quiescence could be induced in 3D alginate hydrogels (Supplementary Figure 2A) that mimic the bone marrow microenvironment, including the presence of RGD peptides (Supplementary Figure 2B), high porosity and low stiffness (Supplementary Figure 2C). hMSCs were expanded in 2D and then cultured with normal medium in 3D alginate-RGD low, where they could interact with RGD motifs in a 3D environment (Figure 2A). After 7 d, hMSCs showed no proliferation as evidenced by almost no EdU-positive cells (Figure 2B).This 3D microenvironment seems, therefore, to promote hMSCs cell cycle arrest. In contrast, hMSCs cultured with starving medium on 3D alginate hydrogels at day 7 showed no cell proliferation (Supplementary Figure 3C). Together, these data indicate that the 3D alginate microenvironment promotes hMSCs cell cycle arrest even in the presence of FBS.

**Figure 2:**
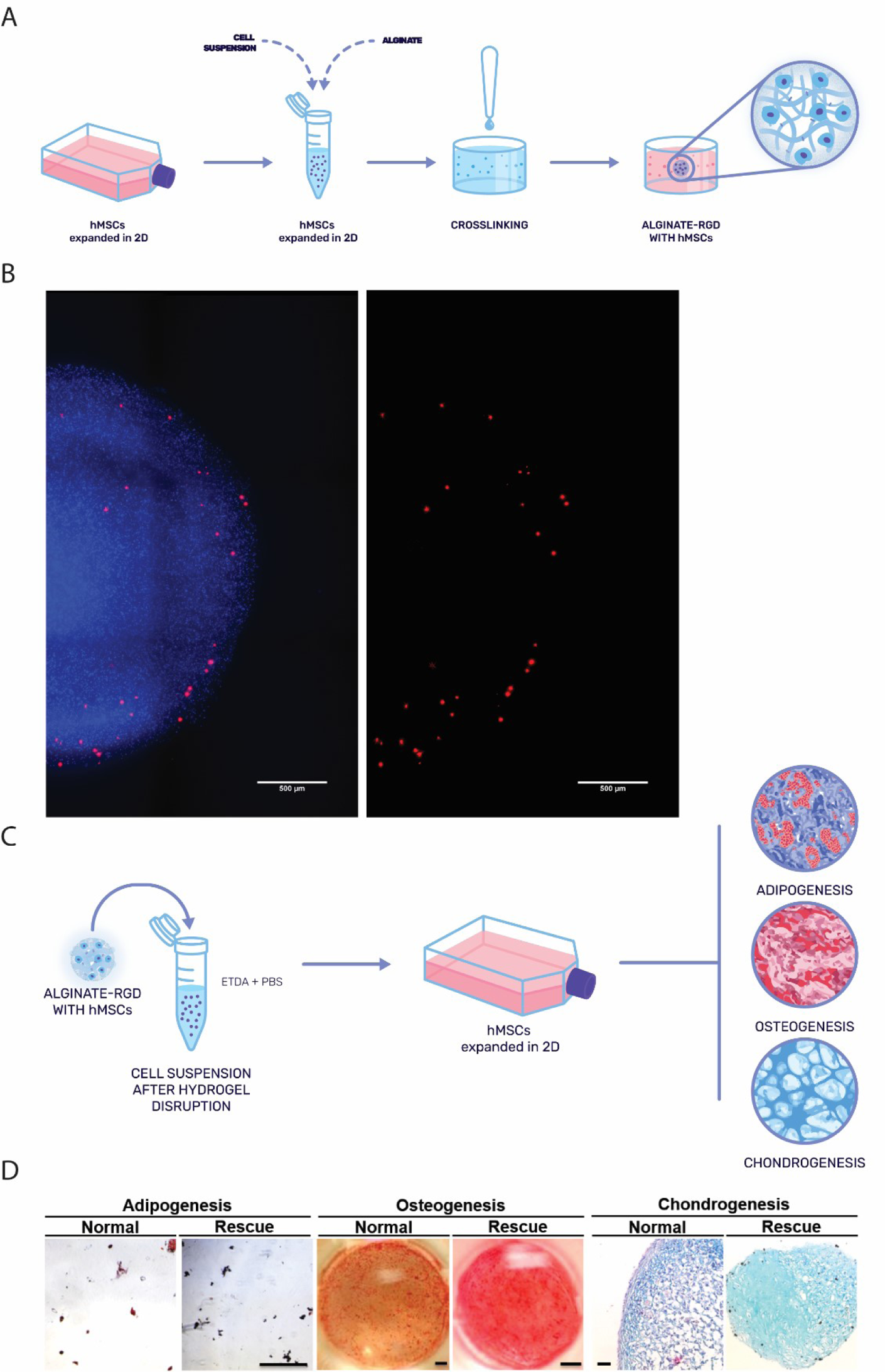
3D alginate-RGD low culture induces hMSCs quiescence. (A) hMSCs encapsulation onto Alginate-RGD low hydrogel experimental setting. Briefly, hMSCs were expanded in 2D, harvested and then mixed with alginate-RGD low. The hydrogels were then cross-linked with CaCl2 and transferred to culture medium. (B) EdU immunofluorescence images of hMSCs cultured in 3D alginate-RGD low hydrogels after 24 hours EdU incubation. In blue nuclei (DAPI), and in red proliferative cells (EdU), n=3; scale bar 500 μm. (C) hMSCs multipotency assessment. Briefly, after culture in alginate-RGD low, hMSCs were incubated in EDTA+PBS to obtain a cell suspension. The cells were then expanded in 2D, and subsequently differentiated into the three classical lineages. (D) After being culture for one week in Normal and Rescue conditions in 3D alginate-RGD low hydrogels, hMSCs were retrieved from the hydrogel, expanded in 2D TCPS and underwent adipogenic, osteogenic and chondrogenic differentiations for 21, 28 and 28 days, respectively. Bright field images show staining for oil red O, alizarin red and alcian blue for adipogenic, osteogenic and chondrogenic differentiations, respectively. n=3; Scale bar: 100 μm.

To determine whether hMSCs cultured on 3D alginate hydrogels maintained their ability to differentiate into the three musculoskeletal classical lineages, we recovered them from the 3D hydrogels, expanded them in 2D (Figure 2C and D) until reaching flask confluency, after which the specific three lineages differentiation protocol was performed. Cells grown in 3D alginate cultured in normal or rescued conditions showed oil red O, alizarin red, and alcian blue staining indicative of adipocytes, osteoblasts, and chondrocytes, respectively.

### 3D alginate-RGD low culture sustains long-term hMSCs quiescence

Having shown that 2D TCPS in starving condition and 3D alginate-RGD low in normal condition induced hMSCs quiescence with different multipotency outcomes, we wondered how these two distinct environments would influence long-term hMSCs quiescence. hMSCs were therefore cultured in 2D TCPS starving and normal conditions and in 3D alginate-RGD low for 28 days. Figure 3A shows how hMSCs metabolic activity changed over time for the three conditions. In normal conditions, hMSCs increased their metabolic activity until day 11, in their proliferation phase, reaching a metabolic activity plateau concomitant with total plate confluence. Metabolic activity did not further increase, probably due to cell-cell contact inhibition hindering cell proliferation [21]. In contrast, in starving condition cells display virtually no metabolic activity after 7 days in culture. This is probably a result of the lack of nutrients, showing that inducing quiescence through starvation is a time-limited process that eventually leads to cell death. In alginate-RGD low, however, hMSCs decreased their metabolic activity until day 11, when they reached a plateau like 2D TCPS normal condition. Despite this plateau being significantly inferior (around 12 times less than 2D TCPS normal condition), it showed that hMSCs survived in 3D over the course of 28 days. In fact, it has been reported that quiescence or cells in G_0_ phase leads to low metabolic activity [22, 23]. This result was supported by live/dead results in which hMSCs cultured in alginate-RGD low were mostly alive (Figure 3B), both in the center or periphery of the hydrogel, while in starving conditions the majority of cells were dead or have vanished with the course of time (Supplementary Figure 3A). During this long-term 3D culture, hMSCs remained mostly quiescent, only having EdU-positive cells in the periphery of the hydrogel (Figure 3C), where cells were no longer exposed to the 3D environment but instead to a more 2D-like environment growing on the surface of the hydrogel. At the periphery, cells adopted a spread shape opposed to a more spherical shape in the center of the hydrogel (Figure 3B).

**Figure 3:**
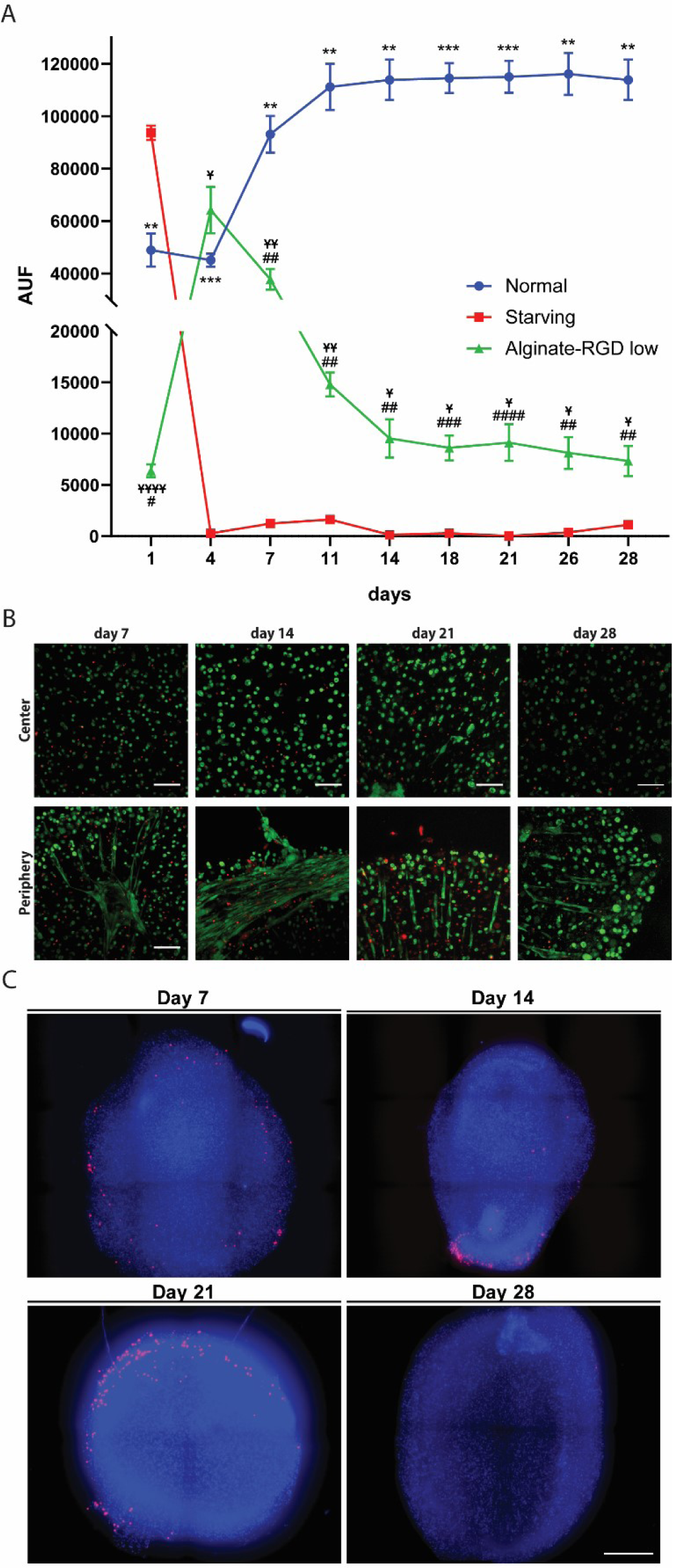
3D alginate-RGD low culture sustains long-term hMSCs quiescence. hMSCs were cultured over a period of 28 days encapsulated in alginate-RGD low. (A) Metabolic activity of hMSCs in 2D Normal and Starving and in 3D Alginate-RGD low. Two-way ANOVA, *, p<0.05; **, p<0.01; ***, p<0.001 when comparing Normal with Starving. #, p<0.05; ##, p<0.01; ###, p<0.001, ####, p<0.0001 when comparing Normal with Alginate-RGD low. Ұ, p<0.05; ҰҰ, p<0.01; ҰҰҰҰ, p<0.0001 when comparing Starving with Alginate-RGD low. (B) Live/dead assay of hMSCs on day 7, 14, 21, and 28. Images were taken in the periphery and center of the hydrogel. Ethidium (in red) stains dead cells while Calcein (in green) stains live cells. Scale bar: 100 µm. (C) Immunostaining for proliferative hMSCs with EdU (in red). Images were taken at day 7, 14, 21 and 28. Scale bar: 500 µm.

This shows that alginate-RGD low allows long-term survival of hMSCs while keeping them in quiescence/G_0_ phase as opposed to starving conditions, which seems only to be able to induce quiescence transiently at the cost of more cell death. This quiescence induction in alginate-RGD low hydrogels seems not to be dependent on a lack of nutrients generated by the hydrogel diffusion dynamics, as alginate-RGD low treated with starving condition showed that the hydrogel cannot support live cells at long-term and ultimately leads to no metabolic activity (Supplementary Figure 3B), no periphery cell proliferation (Supplementary Figure 3C) and overall more cell death (Supplementary Figure 3D).

### hMSCs encapsulated in 3D alginate-RGD low enter in cell cycle arrest and express quiescence genes

Cell cycle regulation is a crucial process for cell survival. The molecular events that occur during this process prevent uncontrollable division and ensure the detection and repair of genetic damages. The main players are cyclins, which form complexes with cyclin-dependent kinase (CDK) and inhibitors of these cyclins [24]. In quiescence, for instances, the cell cycle effectors are inhibited and cells do not progress into the cell cycle, being “arrested” in G_0_ phase. This inhibition can happen through cyclin-dependent kinase inhibitors (CDKNs) and transcription factors [22]. These two types of proteins have a crucial role in preventing cells to undergo proliferation and inducing a quiescence state. At the epigenetic level, other proteins also contribute to the overall quiescence homeostasis, by changes to specific genes through chromatin condensation, allowing or preventing some genes to be accessible to RNA polymerase [23]. To confirm that hMSCs cultured on 3D alginate-RGD hydrogels truly entered quiescence, we quantified the expression of three different genes known to be upregulated in quiescent ASCs [25]: (i) cyclin-dependent kinase inhibitor 1B (p27), a cyclin E and D inhibitor [26]; (ii) forkhead box O3 (FoxO3), a transcription factor that promotes p27 transcription amongst other functions [27]; and (iii) histone-lysine N-methyltransferase (EZH1), an enzyme that has been shown to be essential for hematopoietic stem cell (HSCs) quiescence [28] (Figure 4A). All the three genes were significantly upregulated in hMSCs cultured on 3D alginate-RGD low hydrogels and not in 2D TCPS starving when compared to 2D TCPS normal condition after 6 day. Surprisingly, hMSCs had also significantly higher expression of cyclin E_1_ (marker of late G_1_ phase) and B_1_ (marker of G_2_ and M phase) and a downregulation of cyclin A_2_ (marker of S phase; Supplementary Figure 4).

**Figure 4.**
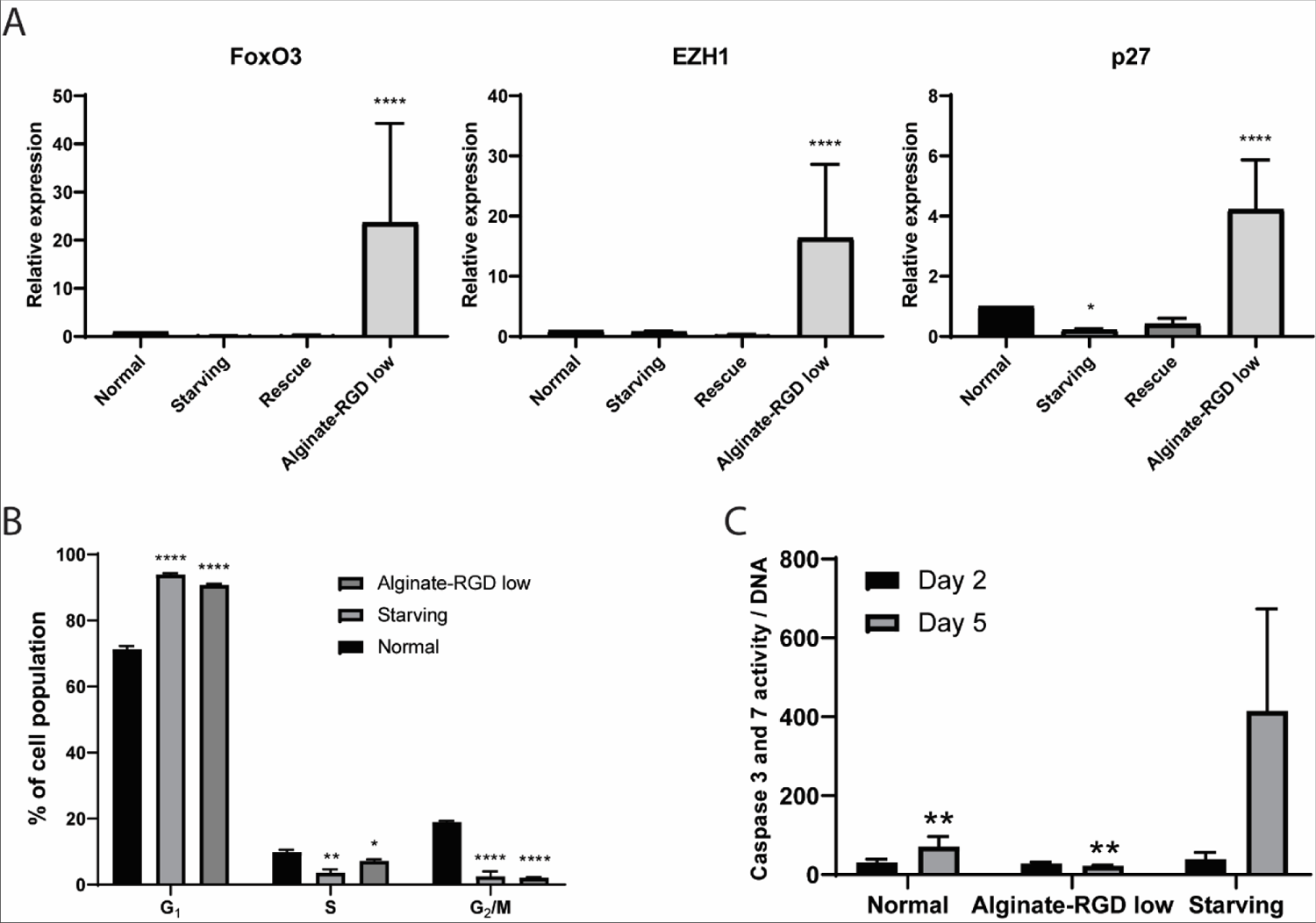
hMSCs encapsulated in 3D alginate-RGD low enter in cell cycle arrest by expressing quiescence genes. (A) gene expression of proteins involved in quiescence after 7 days in Normal, Starving and Rescue 2D and in alginate-RGD low 3D conditions: FoxO3, EZH1 and p27. All values are normalized to Normal 2D condition. (p<0.05, *, p<0.01, **, p<0.001, ***, p<0.0001, ****, n=3) (B) Cell cycle analysis of hMSCs retrieved from Normal and Starving 2D and Alginate-RGD low 3D conditions, after being culture for 7 days (p<0.05, *; p<0.01, **; p<0.0001, **** when comparing to Normal, n=3). (C) caspase 3 and 7 activity of hMSCs cultured in Normal and Starving 2D and Alginate-RGD low 3D conditions after being culture for 2 and 5 days. Multi-comparisons two-way ANOVA was used to compare between Normal, alginate-RGD low and starving conditions (p<0.01, **, between Normal and Alginate-RGD low when compared to Starving n=3).

To better understand why these markers were expressed in the absence of cell proliferation, we analyzed hMSCs cell cycle through propidium iodide double-strand DNA quantification by flow cytometry at day 6 (Figure 4B). G_1_ phase in starving and alginate-RGD low was 1.32 ± 0.01 and 2.74 ± 0.01 times higher than in normal condition, respectively. S phase in normal condition was 2.74 ± 0.57 and 1.38 ± 0.10 times higher than starving and alginate-RGD low, respectively. As for G_2_/M phases, in normal condition cells were 7.47 ± 3.16 and 8.91 ± 0.63 times higher than starving and alginate-RGD low, respectively. These results show that more cells were halted in G_1_ than at S or G_2_/M phases when cultured in 3D alginate-RGD compared to normal 2D TCPS. These results corroborate our previous results with EdU and metabolic activity analysis, showing that both alginate-RGD low and FBS starvation are promoters of cell cycle arrest. When a cell is halted in G_1_ phase and does not resume its cycle, a possible outcome can be apoptosis [29]. We previously observed this effect in the starvation condition in 2D and 3D, which ultimately led to an absence of metabolic activity (Figure 3A and Supplementary Information Figure 3B) and compromised viability (Supplementary Information Figure 3A and 3D). However, if cells enter G_0_ (become quiescent), apoptosis should not be triggered and cells should remain viable until a stimulus induces them to re-enter the cycle. To quantify apoptosis, we have analyzed caspase 3/7 activity in our conditions (Figure 4C). At day 2, there were no significant differences between normal, alginate-RGD low and starving conditions. These two last conditions, however, were not significantly different between each other. We thus concluded that alginate-RGD low promotes hMSCs quiescence, without inducing apoptosis as opposed to starving condition where only a small portion of cells were able to survive upon FBS deprivation. These results are in line with the metabolic activity assay where starving condition reached a very low value of metabolic activity after 11 days of culture.

### Ruling out physicochemical parameters to induce quiescence in 3D hydrogels

Once established that alginate-RGD low was able to induce hMSCs quiescence also at long term and at genetic levels, it became crucial to understand which physicochemical parameters could affect quiescence in 3D. It has been established that the biophysical and biochemical properties of the ASCs niche are fundamental for the maintenance of a quiescent state [2]. Several stem cell niches have been reported to have a hypoxic environment [30] and some researchers have hypothesized that such conditions are ideal for long-term maintenance of stem cells, since in such conditions lesser reactive oxygen species (ROS), responsible for DNA damage, are formed [31, 32]. Some hydrogels, in turn, may recreate a hypoxic environment due to a slower oxygen diffusion. For instance, cells encapsulated in alginate hydrogels with a diameter of 1200 μm had more than 70% of its diameter with a hypoxic core [33]. Since our hydrogels have diameters superior to 3000 μm, we hypothesized that hypoxia could be a driving force behind hMSCs quiescence in our alginate-RGD hydrogels. When cultured in alginate-RGD low, hMSCs became indeed hypoxic (Supplementary Figure 5A) with a brighter fluorescence signal than immortalized MSC (iMSC) cultured in TCPS 2D with 1% (v/v) O_2_ (the brighter, the more hypoxic cells are). However, when hMSCs were cultured in TCPS 2D under hypoxia and normoxia, there were no significant differences between the percentage of cells that underwent proliferation (Supplementary Figure 5B). This led us to conclude that hypoxia is not the main or only factor for alginate-RGD low to drive hMSCs quiescence.

Since lack of nutrients and low oxygen tension cannot be accounted for hMSCs quiescence in alginate-RGD low, we hypothesized that cells became quiescence due to biochemical rather than biophysical cues. Therefore, we cultured hMSCs in a chemically different hydrogel such as collagen. RGD and collagen interact with cells via distinct sets of integrin pairs; namely αvβ3, α5β1 and αIIbβ3 for RGD [34] and α2β1 and α11β1 for collagen [35]. When cultured on top of both alginate-RGD low and collagen hydrogels (Figure 5A), removing thus the potential effects that a 3D environment could introduce to our system, the hMSCs EdU positive population was significantly reduced to less than half when compared to normal condition. Interestingly, even though both materials have different chemistries and interact differently with cells, there were no significant differences in the percentage of EdU positive cells between alginate-RGD low and collagen hydrogels.

**Figure 5.**
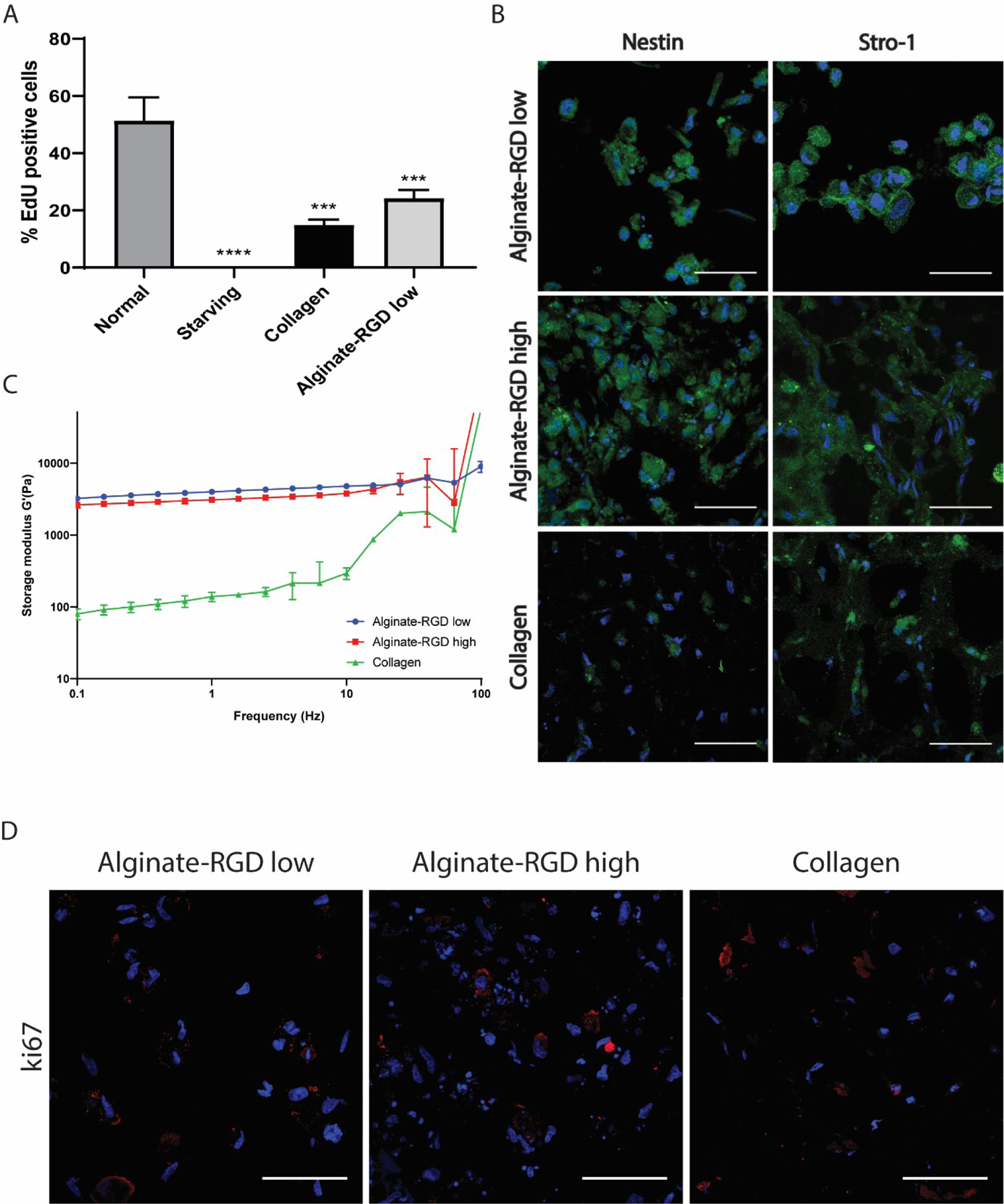
3D soft chemically different hydrogels induce hMSCs quiescence while keeping their stemness. (A) Percentage of EdU positive cells grown for 7 days on top of Normal, Starving (TCPS), collagen and alginate-RGD low hydrogels in 2D. n=3. One-way ANOVA with multiple comparisons (*** p<0.001, **** p<0.0001 to Normal). (B) Nestin and Stro-1 immunofluorescence stainings on hMSCs encapsulated for 7 days in Alginate-RGD low, Alginate-RGD high and Collagen hydrogels in 3D. Nuclei stained in blue (Hoescht). Nestin and Stro-1 in green. n=3, scale bar = 100 μm. (C) Frequency sweeps of alginate-RGD low, alginate-RGD high and collagen were done under the LVR pre-determined for each hydrogel. Storage modulus (G’) in blue for alginate-RGD low, in red for collagen, and green for alginate-RGD high hydrogels. n=2 for alginate-RGD low, n=2 for collagen and n=4 to alginate-RGD high. (D) ki67 immunofluorescence staining on hMSCs encapsulated for 7 days in Alginate-RGD low, Alginate-RGD high and Collagen hydrogels. Nuclei stained in blue (Hoechst) and ki67 in red. n=3; scale bar = 100 μm.

Quiescence in hMSCs has also been correlated with cell shape [6]. Since cell shape is mostly influenced by formation, maturation and clustering of focal adhesions [36–38], we prepared an RGD-modified alginate with high content of the peptide sequence (alginate-RGD high, Supplementary Figure 2B) and compare it to alginate-RGD low, thus varying the amount of formed focal adhesions.

These three different hydrogels, alginate-RGD high, low and collagen, presented all a stiffness below 5 kPa (Figure 5C), which matches the porcine bone marrow G’ values that vary between 5kPa and 7kPa [39]. Despite all the three hydrogels had G’ values statically different at 1 Hz frequency (alginate-GD high lower than alginate-RGD high, p<0.05; collagen lower than both alginate formulations, p<0.0001), we considered that under a certain stiffness threshold all three hydrogels can be considered soft materials (all below 5 kPa at frequencies from 1-10 Hz) that should exert a similar mechanostimuli to cells. hMSCs cultured in all 3D hydrogel conditions showed significantly lower nuclear/cytoplasm yap ratio than 2D TCPS (Supplementary Figure 6A and B) and a lower number of focal adhesions as seen by paxillin and zyxin staining and quantification (Supplementary Figure 6A, C and D). In addition, 3D collagen had significantly higher ratios of paxillin and zyxin expression over cell area than 3D alginate-RGD low and high (Supplementary Figure 6C and D). We also found that in all 3D hydrogel conditions, hMSCs stained positive for nestin and stro-1 (Figure 5B) – markers for hMSCs stemness state i.e. multipotency [40–44]. In fact, in TCPS culture conditions, hMSCs were only positive for STRO-1 and became positive for nestin only when cells were FBS deprived (Supplementary Figure 7). This seems to suggest that when hMSCs become quiescent, either by FBS deprivation or by being cultured in a 3D hydrogel, as shown here by the lack of ki67 positive cells (Figure 5D), they specifically become nestin positive. Hence, regardless of the hydrogel chemical properties, the 3D environment itself seems to allow hMSCs to keep their stemness and quiescence.

Since soft hydrogels induced hMSCs quiescence, we sought to understand whether increasing alginate stiffness (by increasing the total amount of polymer), would also increase hMSCs proliferation (Figure S8). Despite all alginate-RGD low hydrogels presented live cells (Supplementary Figure 8A) and the storage modulus of the hydrogels increased beyond 10 kPa (Supplementary Figure 8B), there were virtually no EdU positive cells on the different alginates with higher stiffness (Supplementary Figure 8C).

### 3D hydrogels induce an alternative pathway for hMSCs quiescence

Virtually all hMSCs encapsulated on 3D hydrogels became quiescent independently of the chemistry or mechanical properties. However, much of the molecular regulation that allows quiescence to happen in 3D is still unknown, and most likely different from that driven by FBS starvation. After having evaluated the effect of potential oxygen deprivation, the effect of chemically different binding motives presented to cells, the amount of these, and the mechanical properties, we assumed that this alternative scenario of quiescence is mainly driven by the dimensionality of the environment. We have therefore focused on a major energy-sensing pathway, the mammalian target of rapamycin (mTOR), which regulates cell growth, proliferation, motility, survival, protein synthesis, autophagy, and transcription. mTOR can form two different complexes: mTOR complex 1 (mTORC1) and mTOR complex 2 (mTORC2) [45, 46]. mTORC1 is composed of mTOR, regulatory-associated protein of mTOR (Raptor), mammalian lethal with SEC13 protein 8 (mLST8) and the non-core components PRAS40 and DEPTOR. This complex functions as a nutrient/energy/redox sensor and controls protein synthesis. mTORC2 is composed of mTOR, rapamycin-insensitive companion of mTOR (RICTOR), mLST8, and mammalian stress-activated protein kinase interacting protein 1 (mSIN1) [45]. mTORC2 has been shown to function as an important regulator of the actin cytoskeleton through its stimulation of F-actin stress fibers, paxillin, RhoA, Rac1, Cdc42, and protein kinase C α (PKCα) [45]. These proteins complexes are therefore involved in many different key aspects of the cell self-regulation, spanning from proliferation to actin reorganization. In fact, mTORC1 has been proposed to be a key protein complex responsible for the transition between G_0_ to G_Alert_ in stem cells, a state where cells are more readily available to re-enter the cell cycle as an exposure to tissue injury [47]. hMSCs encapsulated in 3D Alginate-RGD low, Alginate-RGD high and collagen hydrogels, had no significantly differences in mTOR and Rictor (mTORC2) expression when compared to 2D TCPS normal and starving conditions (Figure 6A, B). Raptor (mTORC1) was only expressed in 2D TCPS normal and starving conditions (Figure 6A, RAPTOR blot). It seems that 2D conditions signal hMSCs to proliferate, through mTOCR1 activity. For starving condition, where the microenvironment does not favor proliferation, hMSCs still express mTORC1 and probably become G_Alert_, i.e. ready to re-enter the cell cycle upon better nutrient conditions. These results are consistent with a major protein involved in multiple cellular processes such as glucose metabolism, apoptosis, cell proliferation, transcription, and cell migration Protein kinase B (also known as Akt). Akt is known to stimulate mTORC1 activity through phosphorylation of tuberous sclerosis complex 2 (TSC2) and PRAS40, both negative regulators of mTOR activity [48]. Here, we show that activated Akt (phosphorylated at T308) has a significant downregulation in the three 3D hydrogels as compared to normal 2D TCPS (Figure 6C), prompting the hypothesis that when hMSCs are cultured in 3D Akt is not activated and does not promote mTORC1 formation and activation, since Akt (P-T308) is considered a reliable biomarker of mTORC1 function [49–52]. Mori et al. [53] have shown that mTORC1 regulates the transcription factor Fork head box O 3a (FoxO3a) through SGK1 kinase. When mTORC1 was inactivated by p18 depletion, cell proliferation was delayed, by phosphorylation of FoxO3a at Ser314. Despite not showing a significant difference, a trend in FoxO3a being more expressed in 3D vs 2D was observed (Figure 6D). The three different hydrogels have no expression of mTORC1 main subunit, Raptor, which in turn results on the lack of FoxO3 inactivation through phosphorylation, hence its presence in the hydrogels. On the other hand, FoxO3 is known to interact with Cyclin-dependent kinase inhibitor 1B (p27Kip1, referred as p27). As it was previously mentioned, p27 has a key role inhibiting cell cycle progression. Its expression and activity is therefore correlated with cell quiescence. We showed that p27 is upregulated in the three hydrogel systems compared to 2D TCPS (Figure 6E). Although not statistically different, a trend where 3D hydrogels seem to have a higher expression of p27 was observed. These results are also in line with gene expression (Figure 4A), as FoxO3a and p27 were upregulated in alginate-RGD low compared to normal 2D TCPS. Finally, we have also looked at retinoblastoma 1 (RB1), which function is to prevent excessive cell growth by inhibiting cell cycle progression until a cell is ready to divide [54, 55]. We found that RB1 is only expressed in 2D starving condition. RB1 seems to be the key protein by which hMSCs become quiescent when starved out of nutrients.

**Figure 6:**
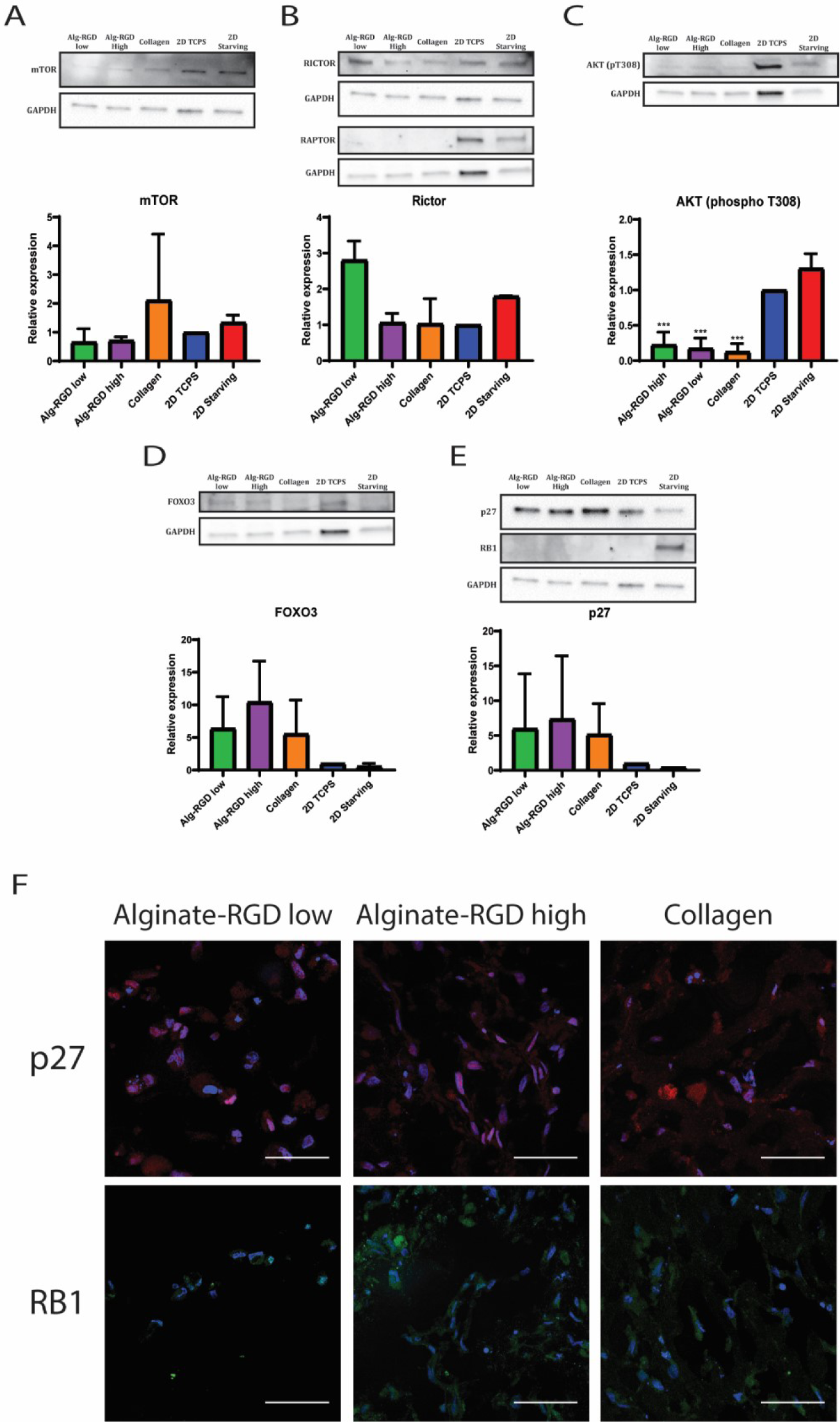
3D hydrogels induce an alternative pathway for hMSCs quiescence. (A), (B), (C), (D) and (E) Protein expression through western blot analysis to hMSCs cultured in Alginate-RGD low, Alginate-RGD high, Collagen, 2D TCPS and 2D Starving. Blot figure and relative expression of each condition to 2D TCPS of (A) mTOR, (B) Rictor and Raptor (without quantification), (C) AKT (phospho T308), (D) Foxo3 and (E) p27 and RB1 (without quantification). n=3, one-way ANOVA, ***, p<0.001. (F) p27 and RB1 immunofluorescence staining on hMSCs encapsulated for 7 days in Alginate-RGD low, Alginate-RGD high and Collagen hydrogels. Nuclei stained in blue (Hoechst). p27 in red and RB1 in green (n=3); scale bar = 100 μm.

These results were also confirmed by immunofluorescence staining of p27 and RB1 (Figure 6F). All three hydrogels stained positive for p27, colocalizing on the cell nuclei where p27 can inactivate cyclins responsible for the cell cycle progression. In contrast, RB1 was negative for all the hydrogels, confirming that hMSCs did not become quiescence due to G1 to S phase blocking through RB1. In contrast, cells cultured in TCPS both under normal and starving conditions did not stain for p27, showing that FBS deprivation quiescence is not regulated by p27 expression, but rather by RB1 (Supplementary Figure 9).

Since our results led us to conclude that soft hydrogels slow hMSCs proliferation rate (2D) or even virtually inhibit it (3D), it was important to understand if stiffness is key to promote quiescence. Stiffer substrate leads to a higher number of mature focal adhesions, which in turn will modulate the cell response. We have hypothesized that hMSCs with lesser number of focal adhesions ultimately would lead to transcript changes towards the same quiescence molecular profile found in 3D. However, when we have inhibited focal adhesions, downstream mechanosensitive pathways or myosin-actin contractibility by treating hMSCs culture in 2D TCPS with Rock inhibitor (Y27632) [56], MRTF inhibitor (CCG-203971) [57], and Blebbistatin [58], respectively, we saw no differences in any of the previous selected proteins (Supplementary Figure 10). Inhibiting two of the major pathways responsible for focal adhesions formation as well as actin stability seems not to have an impact on mTOR (Supplementary Figure 10A), Rictor (Supplementary Figure 10B), Raptor (Supplementary Figure 10C), Akt (Supplementary Figure 10D), p27 (Supplementary Figure 10E) and FoxO3 (Supplementary Figure 10F) expression.

### Alginate-RGD low induces quiescence in vivo

*In vitro* culture of hMSCs encapsulated in alginate-RGD low led hMSCs to become quiescent. However, the higher complexity of an *in vivo* environment poses a challenge for the hydrogel to guarantee the same properties necessary for hMSCs quiescence. We implanted hMSCs encapsulated in alginate-RGD low *in vivo* on a subcutaneous implantation nude rat model (RNU Rat). Alginate-RGD low hydrogels (in red) kept their overall integrity after 3 weeks of implantation, with some fractures due possibly to mechanical stress, chemical degradation, and tissue invasion (Supplementary Figure 11A). 48 hours before euthanizing the animals, an EdU intraperitoneal injection was administered, in order to trace *in vivo* hMSC proliferation. The majority of EdU-positive hMSCs were located in the periphery of the hydrogel and on the host tissue (Figure 7A). Inside the hydrogel, only 5.15 ± 7.58% of the cells were EdU positive (Supplementary Figure 11B). This phenomenon seems to follow a gradient with an increase of EdU-positive hMSCs from the center of the hydrogel towards its periphery (Figure 7B and C).

**Figure 7:**
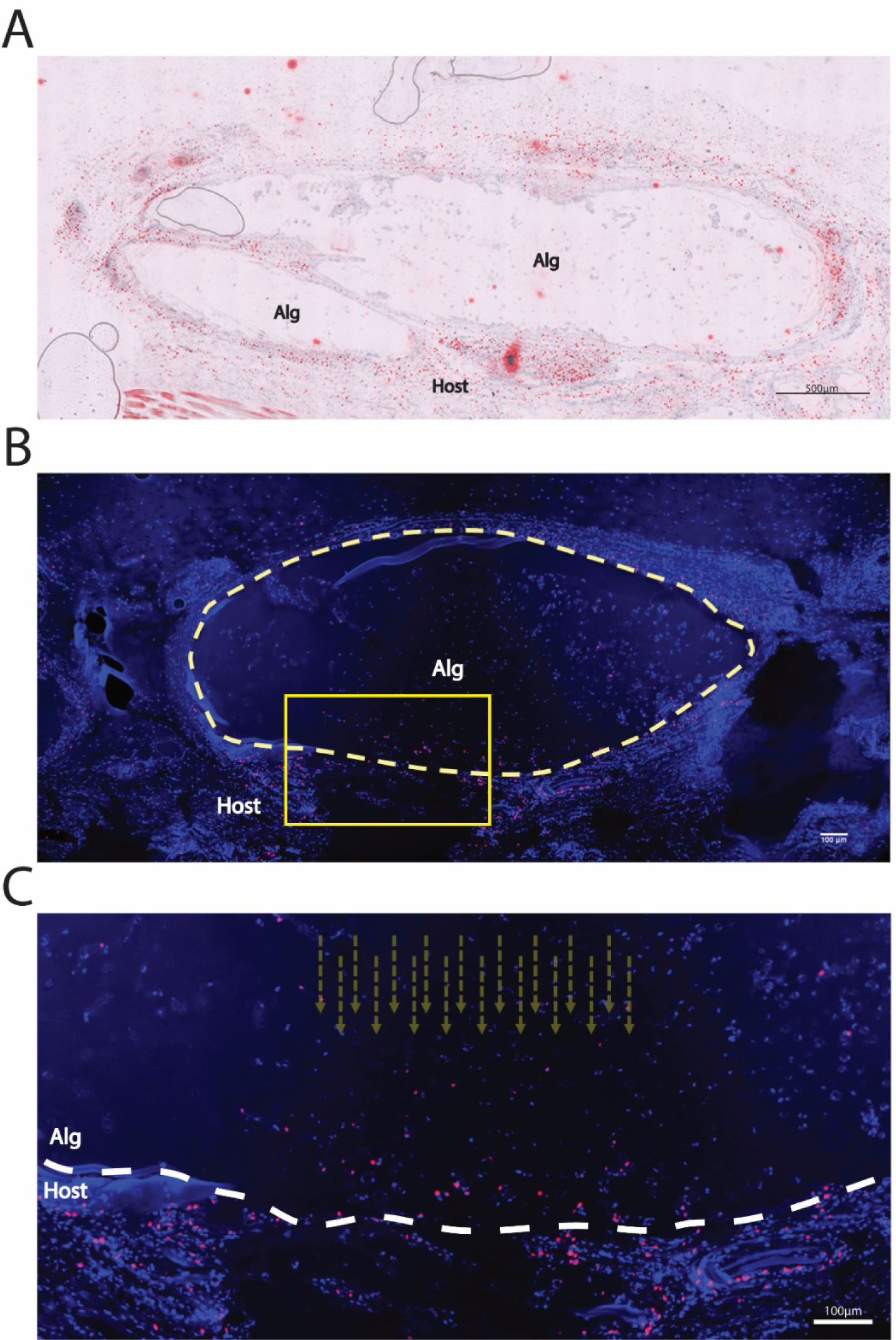
hMSCs encapsulated in Alginate-RGD low in vivo keep their quiescence. EdU immunostaining to hMSCs implanted in pockets formed on the back of rats for 3 weeks. Scale bar: 500 μm. (A) Hematoxylin and EdU stained cut. EdU in red. (B) overview image of EdU positive cells (in red) with a dash line limiting the alginate disc. Nuclei in blue (Hoechst). Scale bar: 100 μm. (C) Magnification of (B) with dash lines showing EdU positive hMSCs gradient. Scale bar: 100 μm. n=8. Alg stands for Alginate-RGD low hydrogel and Host for the host tissue.

These results are in line with the EdU staining shown above, where the only EdU-positive hMSCs were also found in the hydrogel periphery.

## Discussion

Niche intrinsic properties are fundamental to keep ASC quiescence. The depletion of these cells in mammalian tissues has been shown to result in a poor cell turnover and response to injury, ultimately leading to an unhealthy tissue [2, 59]. Here, we have shown that hMSCs proliferate in 2D TCPS surfaces (Young modulus > 1 GPa) until reaching confluency where contact-contact inhibition takes place, which is the opposite of their *in vivo* behavior as also previously reported [60]. When cells were induced to become quiescent through FBS deprivation (starving condition) [20, 61], they showed virtually no proliferation with the ability to resume their cell cycle when FBS was again introduced five days after (Figure 1 B and C). Despite being multipotent and remaining undifferentiated after this starvation treatment, showing differentiation towards the three classical hMSCs lineages, the surviving cells surprisingly showed less osteogenic and chondrogenic differentiation (Figure 1E). Serum starvation is often used as a method for cell synchronization, which facilitates cell (re)programming [62] and, therefore, differentiation. We would then expect that starved hMSCs would perform better differentiations with deposition of larger amount of characteristic ECM.

For most of ASC, quiescence and multipontecy can be often assessed by specific markers expressed on the cell surface. For instances, HSC can be identified through c-Kit^+^Sca-1^+^Lin^-^ Tie2^+^, c-Kit^+^Sca-1^-^Lin^-^CD150^+^CD48^-^CD34^-^, and c-Kit^+^Sca-1^+^Lin^-^CD48^-^CD150^+^, while MuSC through SMC2.6^+^CD45^-^, M-cadherin^+^, and c-met^+^ [22]. For hMSCs, it is still not fully understood whether they can become quiescent or not, hence the difficulty to identify markers for a quiescent state. However, some researchers have proposed that both Nestin^+^ [43] and PDGFRα^+^Sca-1^+^CD45^-^TER119^-^ [53] are markers for quiescent MSC in *in vivo* mouse models. In fact, in both these studies, these MSC populations were not only in a resting state, but were also able individually to originate hematopoietic niche cells when transplanted *in vivo* for more than one sequential transplantation. Interestingly, when Nestin^+^ MSC were cultured in TCPS, they also started proliferating and lost their Nestin expression with passaging. In our culture conditions, when hMSCs were cultured with FBS on TCPS, they also did not express Nestin. However, when deprived of FBS, some hMSCs became Nestin^+^, which suggests that Nestin is indeed associated with quiescence (Supplementary Figure 2).

Despite these results, FBS deprivation does not recapitulate the *in vivo* microenvironment. *In vivo*, hMSCs are often surrounded by other cell types in a hypoxic niche, in a low stiff rich ECM microenvironment [22, 63, 64]. In order to better mimic these conditions, we have chosen to culture hMSCs in a 3D hydrogel where we could meet some of the native hMSCs niche characteristics. Alginate allowed us to create a hypoxic environment, while providing biological (RGD peptides) and biophysical (low stiffness) cues (Figure 2A). hMSCs showed no proliferation in alginate-RGD low, which has been shown to happen by others in non-dynamic hydrogels like ours [65–67]. After being in 3D culture for 5 days, cells were able to resume their cell cycle, shown by re-population of 2D TCPS flasks until full confluency. In fact, Maia et al. [66] has also shown that hMSCs encapsulated in alginate-RGD hydrogels did no proliferate, reduced their metabolic activity and kept their DNA content, regardless of the cell density. When retrieved from these hydrogels, cells were able to proliferate again in TCPS 2D. Surprisingly, when cultured in our alginate-RGD low hydrogels, hMSCs differentiated into the three hMSCs classical lineages without showing a reduction in their multipotency as opposed to starving condition (Figure 2D). Quiescence in hMSCs can then be induced through culture on a FBS deprived medium (starving) or in 3D in an alginate-RGD low hydrogel. We confirmed it by analyzing hMSCs cell cycle. In both alginate-RGD low and starving conditions, more than 90% of cells analyzed were at G_1_ phase, while S and G_2_/M phases had significant lower values than normal condition (Figure 4B).

Despite both conditions led to quiescence, it was worthwhile exploring the differences thus far noticed between them. The first striking difference was the long-term cell viability in alginate described here. Cultured in alginate-RGD low, hMSCs were able to survive for at least 28 days, keeping a low metabolic activity (Figure 3A). In starving condition, after 4 days in culture, the metabolic activity was nearly zero and almost no viable cells were detected right after (Supplementary Figure 4). These results were further validated by detecting more caspase 3/7 activity in starving condition when comparing to Alginate-RGD low hydrogels (Figure 4C). This indicates that, when starved, hMSCs undergo senescence at higher extent than cultured in 3D with FBS. This shows that the intrinsic properties of alginate-RGD low allow hMSCs to become not only quiescent, but to keep it in such a state for a long period of time. This phenomenon could be explained by the fact that alginate-RGD low hydrogel cultures induced an upregulation of FoxO3, EZH1 and p27 genes expression (Figure 4A). These results are in line with what has been described for other ASC, including HSCs, MuSCs and HFSCs. In a comprehensive review from Cheung et al. [68] about the molecular mechanisms that regulate stem cells, these three genes are described to be key proteins, amongst others, to keep ASC in quiescence. In fact, it has been shown that knocking-down FoxO3 in HSCs lead to a defective maintenance of its quiescence pool [69], while in MuSC its depletion leads to self-renew impairment [70]. EZH1, which is responsible for chromatin modifications associated with quiescence maintenance, was required for HSC quiescence and to prevent their senescence [28]. p27, a CDK inhibitor (CKI), was also upregulated in several ASC. In HSC, p27 has a synergetic interaction with p57, controlling the nuclear transportation of HSC70–cyclin D1 complex, and therefore regulating cell cycle re-entry [71]. In muscle tissue, p27 has been used as detection method for quiescence MuSC [72]. For hMSCs, Rumman et al. have shown that when cultured in polyacrylamide (PAA) 2D hydrogels with a stiffness of 0.6 kPa, the cells expressed p27 and p21 genes and became quiescent with low percentage of ki67 and bromo-deoxyuridine (BrdU) positive cells [73].

Surprisingly, in our study alginate-RGD low hydrogels are not the only ones we have reported to induce hMSCs quiescence. Alginate-RGD high, which has more RGD units than alginate-RGD low, providing more integrin connection points, and collagen hydrogels, which interact with cells through different integrins than those interacting with RGD [34, 35], also showed virtually no proliferation (Figure 5D). While some cells became nestin^+^ in starving condition, for all types of 3D hydrogels virtually all hMSCs were positive for nestin (Figure 5B), showing that 3D hydrogel culture seems to revert hMSCs towards a quiescence state. We have not only explored biochemical changes, but also looked into biophysical changes. We have shown that hMSCs grown in surfaces with lower mechanical properties have roughly halved their EdU-positive population in 2D (Figure 5A). However, by increasing the alginate content on our alginate-RGD low hydrogels, we have seen no changes in the EdU-positive population, i.e. no changes in proliferation (Supplementary Figure 8). Of course, the observed changes in storage modulus might not be enough to increase hMSCs proliferation, as changes from alginate-RGD low 2D surfaces to TCPS stiffness cannot compare to the maximum measured storage modulus of 10 kPa approximately (Supplementary Figure 8B).

In view of the latest data on ASC quiescence, some researchers have been proposing that quiescence can in fact be distinguished by separate states that vary in depth, with the limits of the spectrum being “deeper” quiescence and quiescence “alert” [22, 75]. Upon an injury in the tissue, ASC become quiescence “alert”, and ready to re-enter the cell cycle to contribute for tissue regeneration. This state is characterized by a propensity for proliferation and differentiation and by the expression and activity of mTORC1. In contrast, in “deep” quiescence cells have less metabolism, which is a result of lack of mTORC1 expression. ASCs switch between these two states, reverting to “deep” quiescence when the tissue regeneration is complete [22]. In our results, hMSCs cultured in all hydrogels systems are mTORC1^-^, suggesting that they are in a “deep” quiescence state kept by FoxO3 and p27 expression (Figure 6). The same does not happen when cells are FBS deprived. Starved hMSCs still expressed mTORC1 and had no upregulation of FoxO3 or p27, but kept their quiescence status by expressing RB1 protein, suggesting that their quiescence is closer to a state similar to quiescence “alert”. In fact, it has been shown that hMSCs cultured as spheroids, grown on top of chitosan films, are also quiescent, become senescent and are mTORC1^+^ like our starving condition. When the spheroids 3D culture was combined with the mTORC1 inhibitor rapamycin treatment, peripheral apoptosis and dead cell attachment were prevented, while increasing pluripotent gene expression [76]. It is quite interesting that the 3D system here explored were somehow able to lead hMSCs to inhibit mTORC1 expression, which in turn might have saved them from senescence.

Finally, our results show evidence that hMSCs encapsulated in alginate-RGD low still keep their quiescence status *in vivo*. Proliferative cells seem to be reserved to the periphery of the hydrogel, as an outwards gradient could be detected (Figure 7). In fact, cells in the hydrogel periphery are more exposed to nutrients, as well as other biophysical and biochemical cues (soluble or non-soluble) from the host tissue that could induce proliferation. The wound site formed by the hydrogel implantation is known to be rich in several factors including cytokines that promote cells to proliferate [77].

This study is a step forward towards understanding better quiescence in hMSCs. While most of the information gathered in literature explores other ASCs, little is known about hMSCs mechanisms to keep a pool of quiescence stem cells throughout mammalian lifespan. Ferreira et al. [78] had already shown that ECM is fundamental for quiescence maintenance, by exploring the role of a glycoprotein-rich pericellular matrix on inducing hMSCs quiescence when cultured at a high cell density encapsulated in thiol-modified hyaluronic acid (S-HA) and poly(ethylene glycol) diacrylate (S-HA-PEGDA) hydrogels. Here, we propose that hMSCs quiescence has a distinct molecular signature when cultured in 2D versus 3D. While in 3D we believe that mTORC is inactivated through an inactivation of Akt signaling, leading to an activation of FoxO3 and, consequently of p27 gene, in 2D FBS deprived cultures, Akt is not inactivated, therefore failing to inactivate mTORC1. Instead, cells in FBS starvation depend upon RB1 to prevent cell cycle re-entry (Figure 8). More in-depth molecular analysis will have to be conducted in order to unravel the full molecular signatures behind “deep” quiescence and quiescence “alert”of hMSCs, and ultimately validate the existence of a clear quiescence state *in vivo* for hMSCs populations.

**Figure 8:**
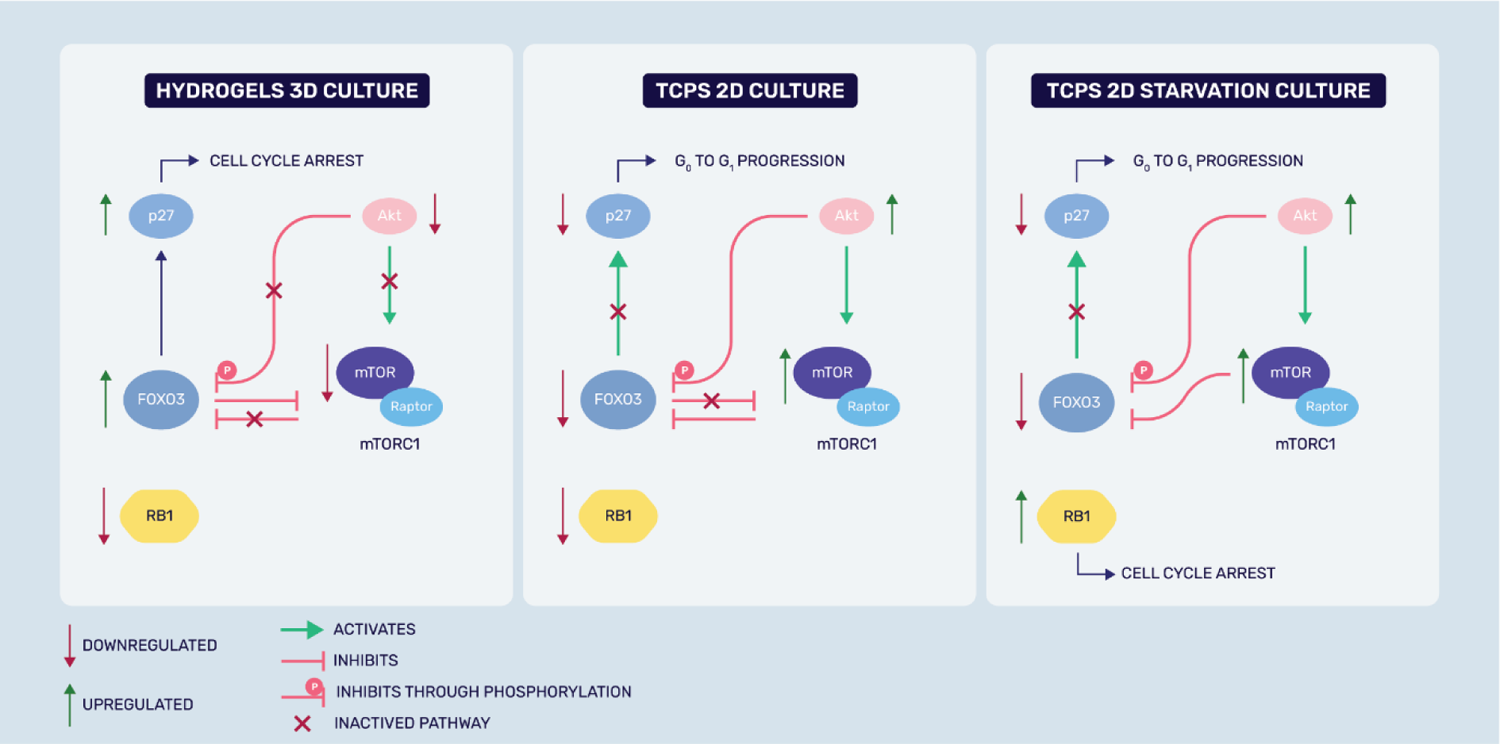
Proposed model for hMSCs quiescence molecular signature in TCPS 2D starvation versus hydrogels 3D culture systems. When hMSCs are culture in 2D TCPS with FBS (normal culture), they express activated Akt which can activate mTORC1. mTORC1 inactivates FoxO3 which can also lead to p27 inactivation. In hydrogels 3D, on contrary, Akt is downregulated, which can also lead to mTOCR1 inactivation. In turn, this inactivation will cause the opposite effect of TCPS 2D culture molecular pathway, with FoxO3 and p27 upregulation. For 2D starvation culture, hMSCs fail to inactivate Akt energy sensory pathway, which potentially leads to keeping mTORC1 active and therefore no FoxO3 or p27 upregulation. Instead, hMSCs quiescence is kept by RB1 protein, which seems not to be involved in hydrogels 3D culture.

## Conclusion

Our new findings described here pave the way towards understanding how quiescence can be induced and maintained in hMSCs when cultured in 3D. This quiescence state in 3D, molecularly distinct from 2D quiescence, shows that dimensionality is key on changing cell’s phenotype. As hMSCs are a useful tool for regenerative medicine therapies and have been combined with biomaterials, a deeper understanding on how these cells behave in 3D could prove useful for future cell-based regenerative medicine strategies.

## Acknowledgments

We are grateful to the European Research Council starting grant “Cell Hybridge” for financial support under the Horizon2020 framework program (Grant #637308). Some of the materials used in this work were provided by the Texas A&M Health Science Center College of Medicine Institute for Regenerative Medicine at Scott & White through a grant from NCRR of the NIH (Grant #P40RR017447). We also would like to express our gratitude to Guilherme Gomes, who kindly helped us on the illustrations on this manuscript, to Andrea Calore for helping with rheological data, as well as to David Koper, Marloes Peters and Rick Claessen who helped us performing part of the *in vivo* studies in this work.

## Supplementary information

**Supplementary figure S1.**
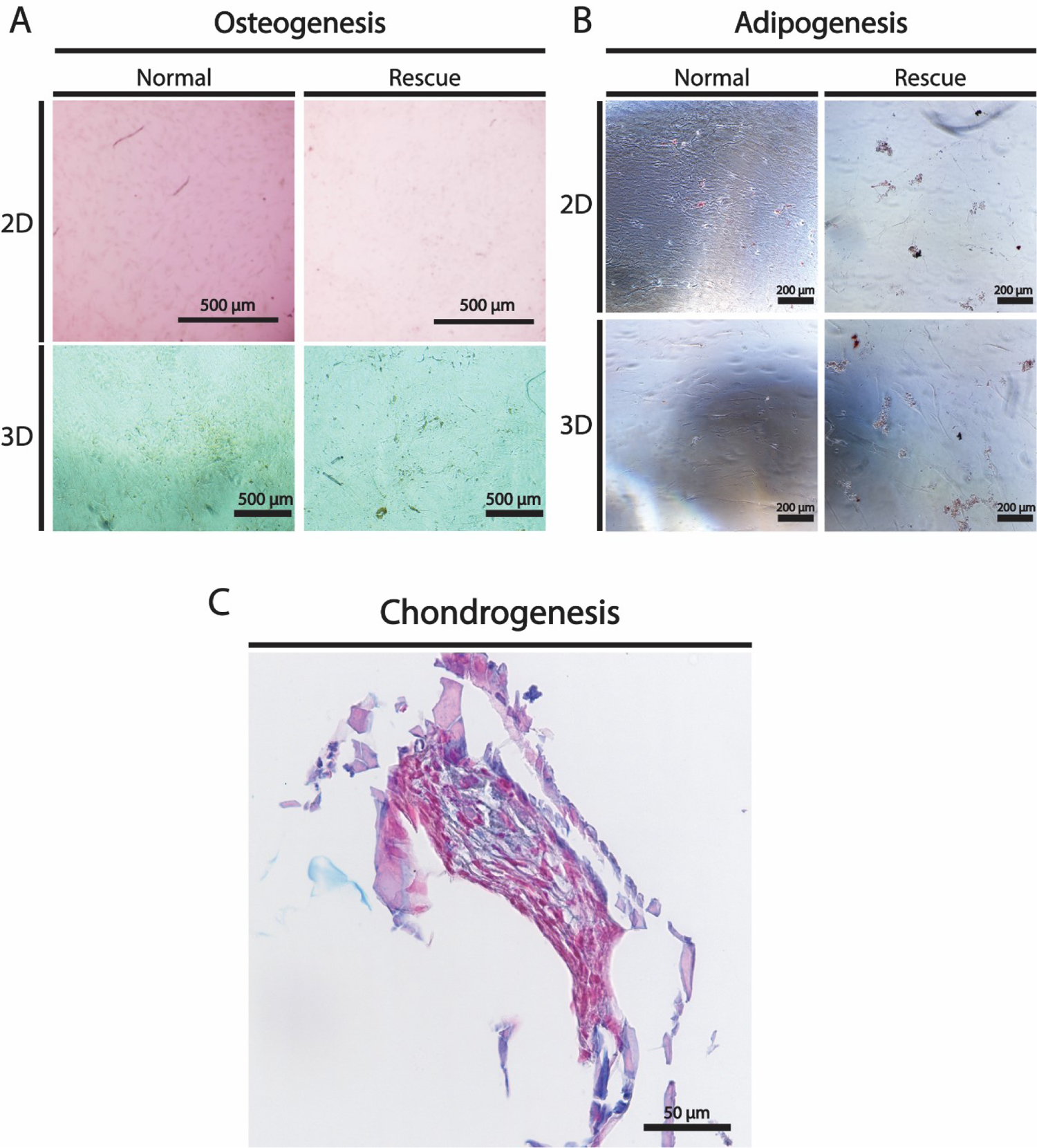
2D and 3D differentiation controls. After being culture for one week in Normal and Rescue conditions in 2D and 3D alginate-RGD low hydrogels, hMSCs were retrieved from the hydrogel, expanded in 2D TCPS and were cultured in basal medium for the same duration of adipogenic (A) and osteogenic (B) differentiations, as controls, for 21 and 28 days, respectively. For chondrogenic differentiation, only normal 2D was cultured under basal medium for controls purposes. Bright field images show that staining for oil red O, alizarin red and alcian blue for adipogenic, osteogenic and chondrogenic differentiations, respectively were negative, meaning that hMSCs cultured in TCPS did not commit to any classical hMSCs cell lineage. n=3.

**Supplementary figure 2.**
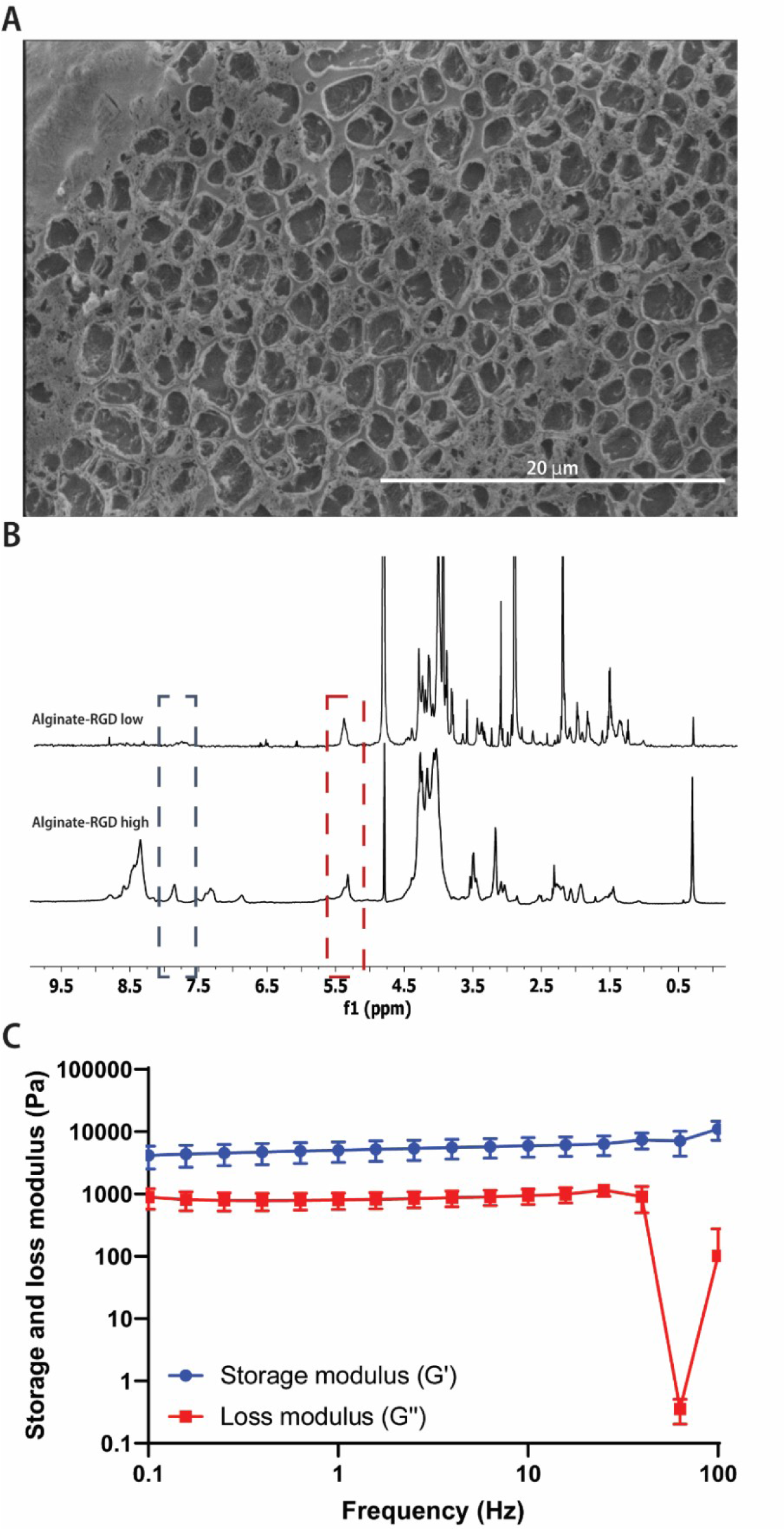
Alginate-RGD low characterization. Alginate-RGD has been analyzed by (A) CryoSEM, (B) NMR and (D) rheology. (A) cryoSEM image of alginate-RGD low shows the material microporosity. (B) H1-NMR spectra of Alginate-RGD low (top) and high (bottom) in sodium acetate-d3 at pH 3. The intensity of the spectra are normalised to the peak corresponding to the anomeric proton of guluronic block (red box) at 5.35 ppm. Blue box shows characteristic peaks of RGD sequence (amine groups) at 7.6-7.8 ppm. From the spectra, it is possible to see that alginate-RGD high has a higher number of RGD sequences attached to the alginate backbone (D) Frequency sweeps of alginate-RGD low done under the pre-determined LVR. Storage modulus (G’) in blue, always under 5 kPa and loss modulus (G’’) in red. n=3.

**Supplementary figure 3.**
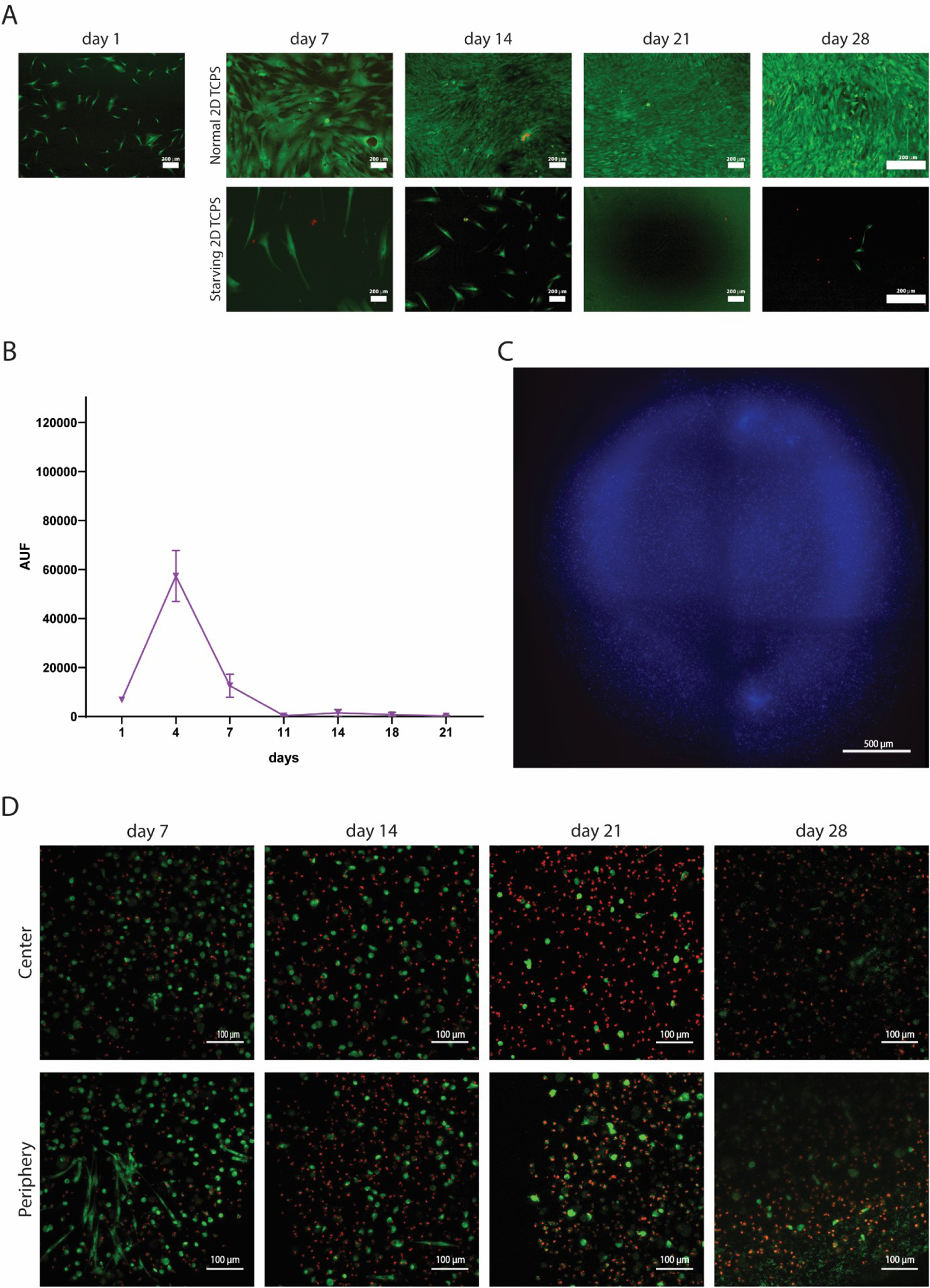
Long-term hMSCs survival. (A) Live/dead assay to normal and starving in 2D TCPS shows that hMSCs did not survive and vanish in the course of time. When not deprived from FBS, they thrive, proliferating until confluency. Scale bar: 200 µm; n=3 (B) hMSCs were cultured over a period of 28 days encapsulated in alginate-RGD low, treated for starving condition. After 11 days there is virtually no metabolic activity (C) Immunostaining for proliferative hMSCs with DAPI (in blue) and EdU (in red) at day 7 showing virtually no EdU^+^ cells. Scale bar: 500 µm. (D) Live/dead assay of hMSCs on day 7, 14, 21 and 28 shows a decrease in cell viability with increasing prevalence of dead cells. Images were taken in the periphery and center of the hydrogel. Ethidium (in red) stains dead cells while Calcein (in green) stains live cells. Scale bar: 100 µm.

**Supplementary figure 4.**
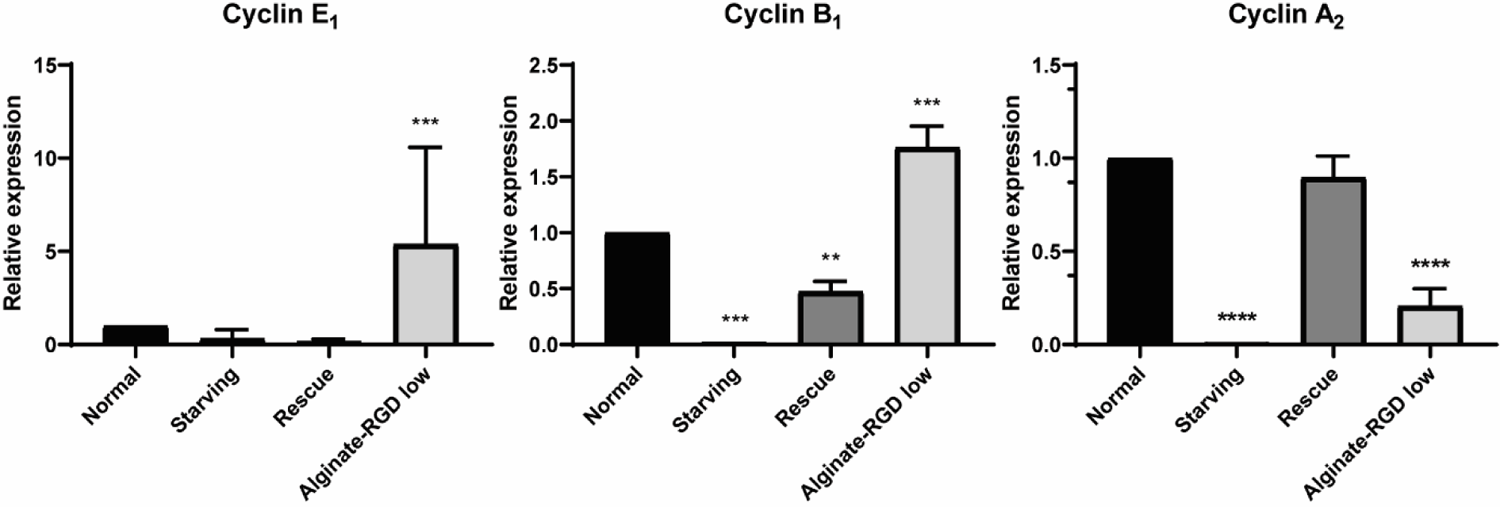
Cyclin gene expression. Gene expression of proteins activated in cycling cells: cyclin E1, cyclin B1 and cyclin A2 after 7 days in Normal, Starving and Rescue 2D and in alginate-RGD low 3D conditions. Alginate-RGD low shows an upregulation of cyclin E_1_ and B_1_. Cyclin A_2_ is, however, downregulated. All values are normalized to Normal 2D condition. (p<0.01, **, p<0.001, ***, p<0.0001, ****, n=3)

**Supplementary figure 5.**
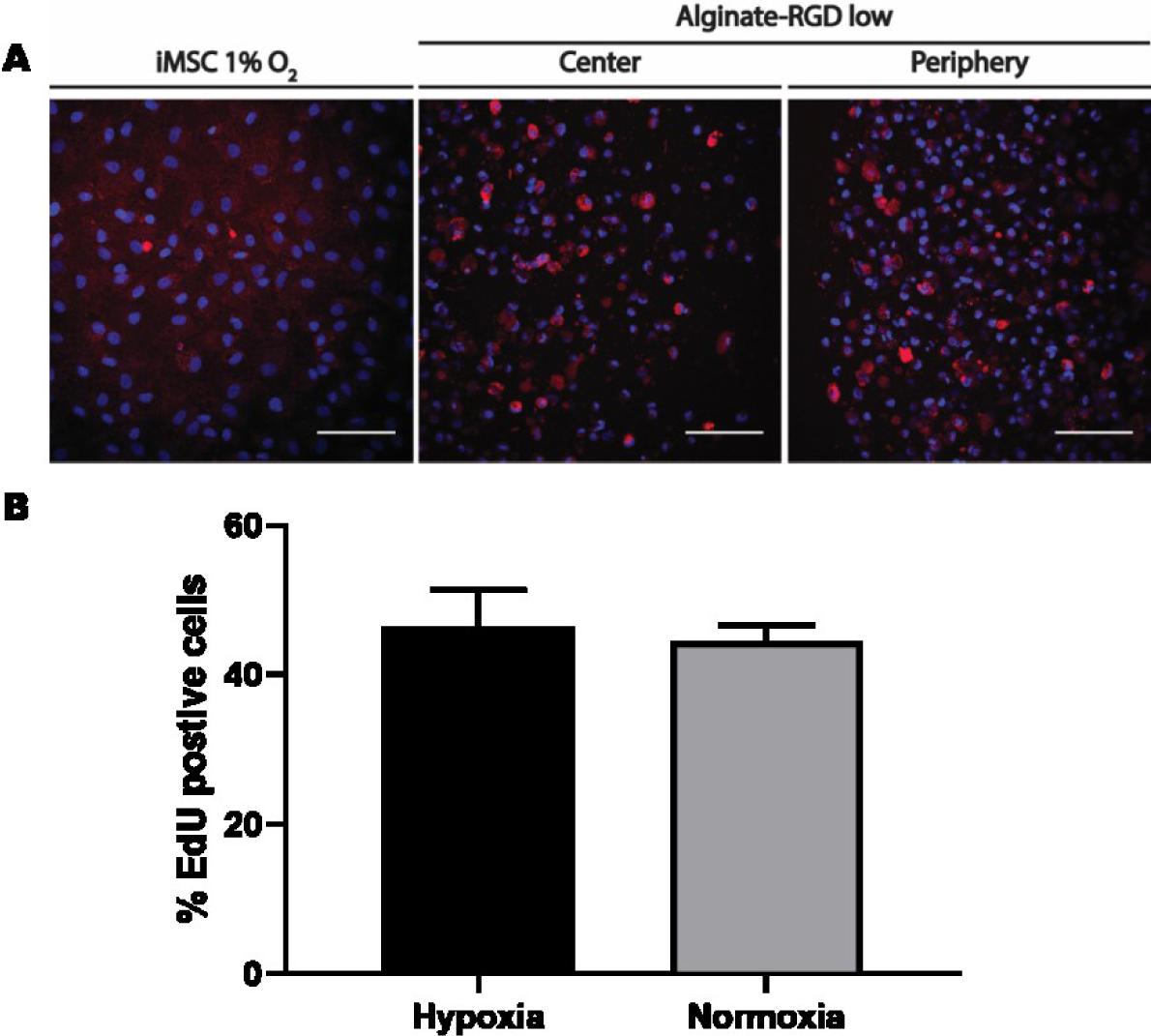
hMSCs hypoxia assessment. (A) After a week, iMSCs cultured in 2D TCPS in hypoxia (control) and hMSCs cultured in 3D alginate-RGD low in normoxia (both center and periphery) stained positive for hypoxia with Image-iT™ Red Hypoxia Reagent (in red) and Hoescht (in blue). Scale bar: 100μm. (B) After a week in culture in 2D TCPS, 24 hours before fixing, hMSCs in hypoxia or normoxia were treated with EdU to assess the percentage of cells that underwent proliferation. There were no statistical differences between both conditions, which shows that hypoxia did not induce quiescence in 2D TCPS.

**Supplementary figure 6.**
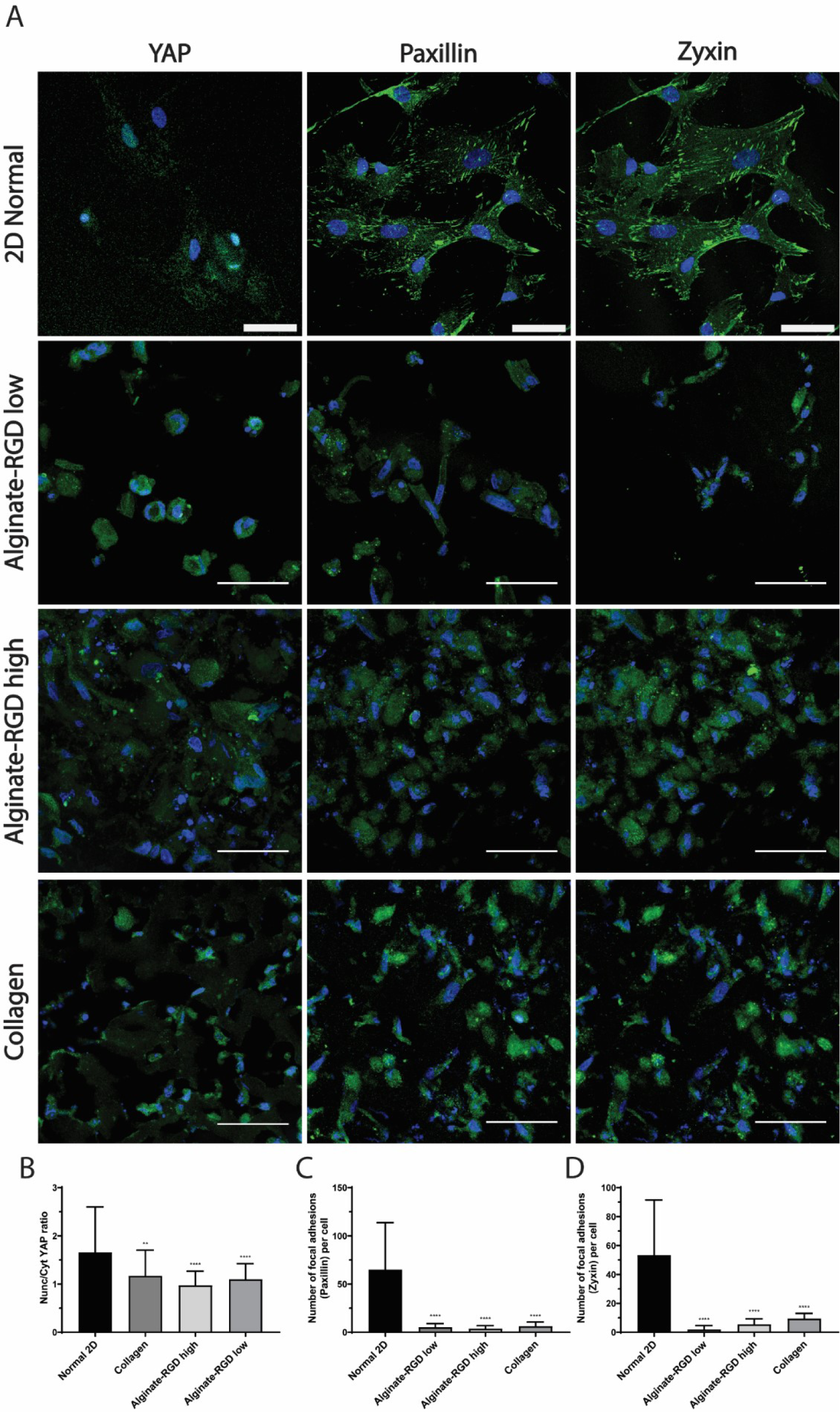
hMSCs 3D mechanosensing. (A) Yap, Paxilllin and Zyxin immunofluorescence staining on hMSCs cultured in 2D TCPS normal and encapsulated in 3D in Alginate-RGD low, Alginate-RGD high and Collagen hydrogels. Nuclei stained in blue (Hoechst). Yap, Paxilllin and Zyxin in green. n=3, scale bar = 50 μm. (B) Nuclei/cytoplasm fluorescence intensity quantification of Yap staining for 2D TCPS, Collagen and Alginate-RGD low and high hydrogels in 3D. (C) and (D) Quantification of number of paxillin positive dots (C) and focal adhesions (D) per cell for Normal 2D, Collagen, Alginate-RGD low and high, showing that when cells are cultured in 2D they experience a stiffer surface that translates in more focal adhesions than any hydrogel studied hereAll graphs have n=20 cells. One-way ANOVA multiple comparisons. ** p<0.01, *** p<0.001, **** p<0.0001.

**Supplementary figure 7.**
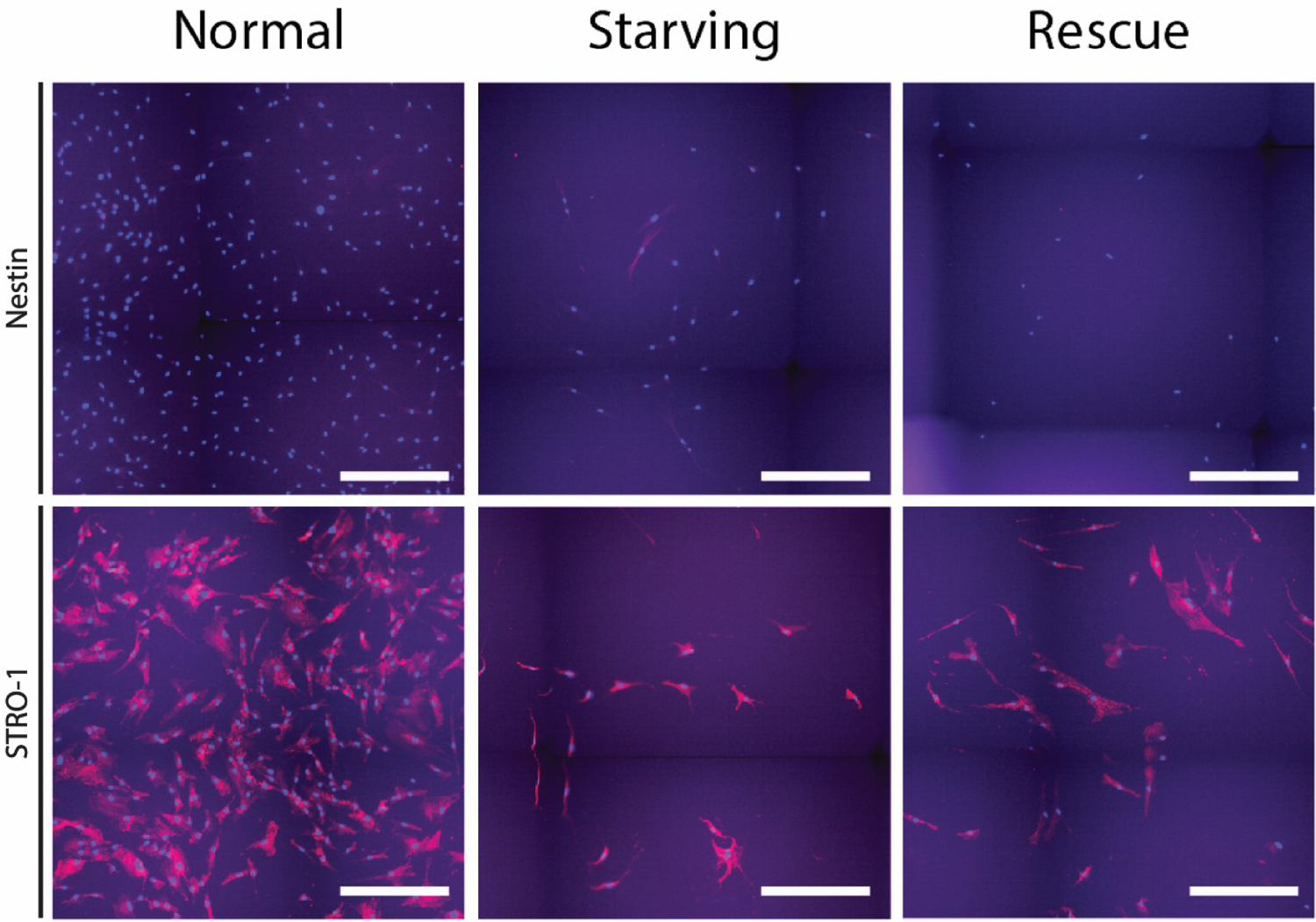
2D hMSCS stemness. hMSCs cultured in TCPS 2D in normal, starving and rescue condition stained with anti-Nestin and anti-STRO-1 with Alexa fluor 568. Stro-1 was expressed regardless of the condition. However, nestin, a potential marker for quiescence, was only expressed transiently in starving condition and only in some cells. Scale bar: 500 µm.

**Supplementary figure 8.**
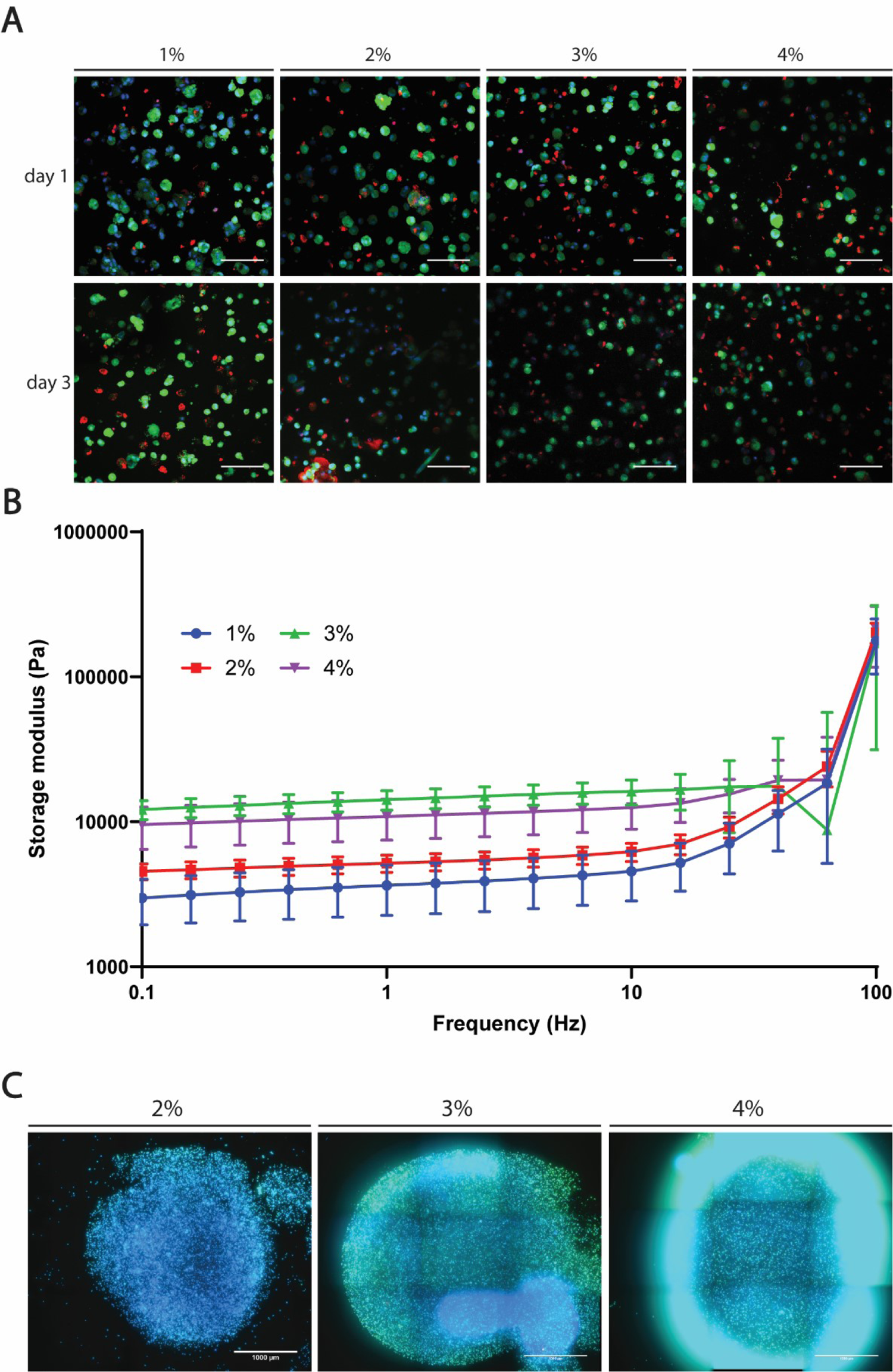
Stiffness influence on quiescence. (A) Live/dead images of hMSCs encapsulated in 1%, 2%, 3% and 4% Alginate-RGD low at day 1 and 3 show that cells are alive in stiffer materials and over time. In green live cells (calcein AM) and in red dead cells (Ethidium homodimer), n=1; scale bar 100 μm. (B) Storage modulus acquired by frequency sweep on a rheometer for Alginate blank at 1%, 2%, 3% and 4% show an increase in stiffness that plateaus on 3%, without an increase after 4%; n=5. (C) EdU immunofluorescence images of hMSCs encapsulated in 2%, 3% and 4% Alginate-RGD low at day 6 after 24 hours EdU incubation showed no EdU^+^ cells, regardless of the material. In blue nuclei (Hoechst), in green cell bodies (Syto14) and in red proliferative cells (red), n=3; scale bar 1000 μm.

**Supplementary figure 9.**
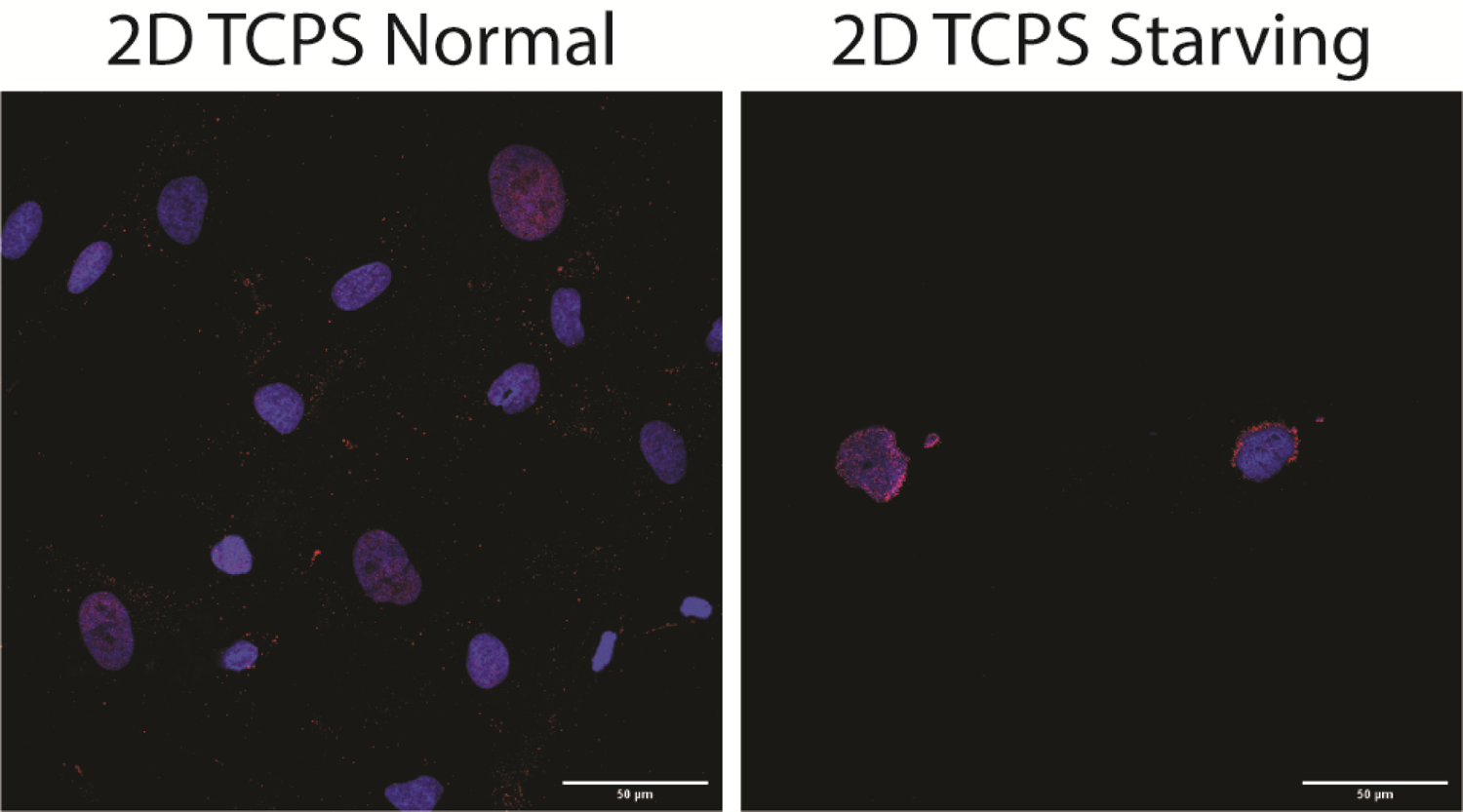
p27 immunostaining in 2D. hMSCs cultured in 2D TCPS normal and starving conditions were stained for p27 at day 7. Both conditions show a barely any p27 staining, indicating that cells grown or deprived with FBS on a 2D surface do not become quiescence through p27 activation. In blue nuclei (Hoechst) and in red p27, n=3; scale bar 50 μm.

**Supplementary figure 10.**
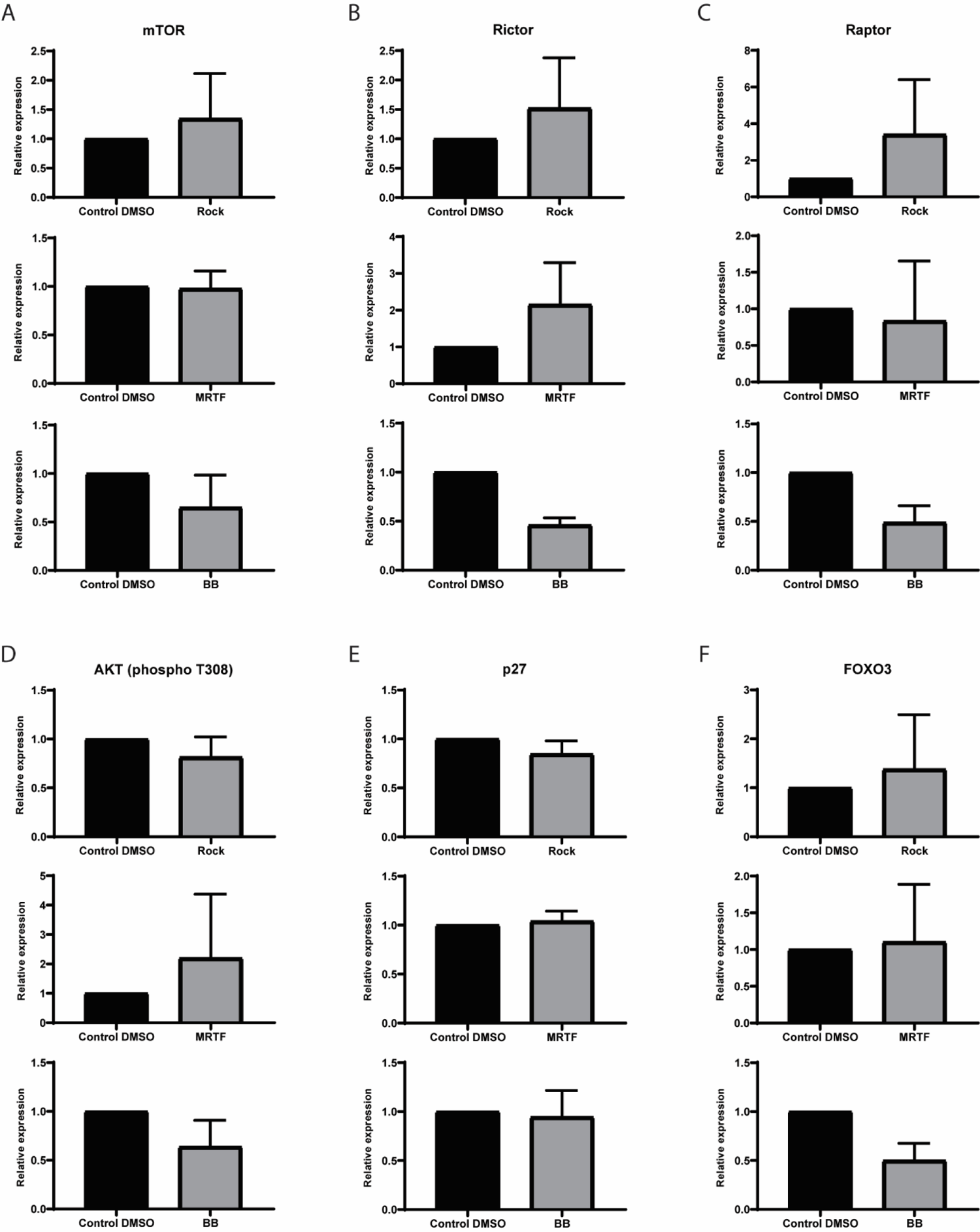
Mechanosensing pathways inhibition. Protein expression through western blot analysis to hMSCs culture in 2D for 7 days and treated with control DMSO, ROCK inhibitor (ROCK), MRTF inhibitor (MRTF) and Blebbistatin (BB). Relative expression of each condition to Control DMSO of (A) mTOR, (B) Rictor, (C) Raptor, (D) AKT (phospho T308), (E) p27 and (E) Foxo3 showed no significant differences between controls and inhibitions. Inhibiting the mechanossensing pathways seemed to not induce quiescence in hMSCs, which did not replicate the conditions that led hMSCs to become quiescence in 3D as it was hypothesized. n=3.

**Supplementary figure 11.**
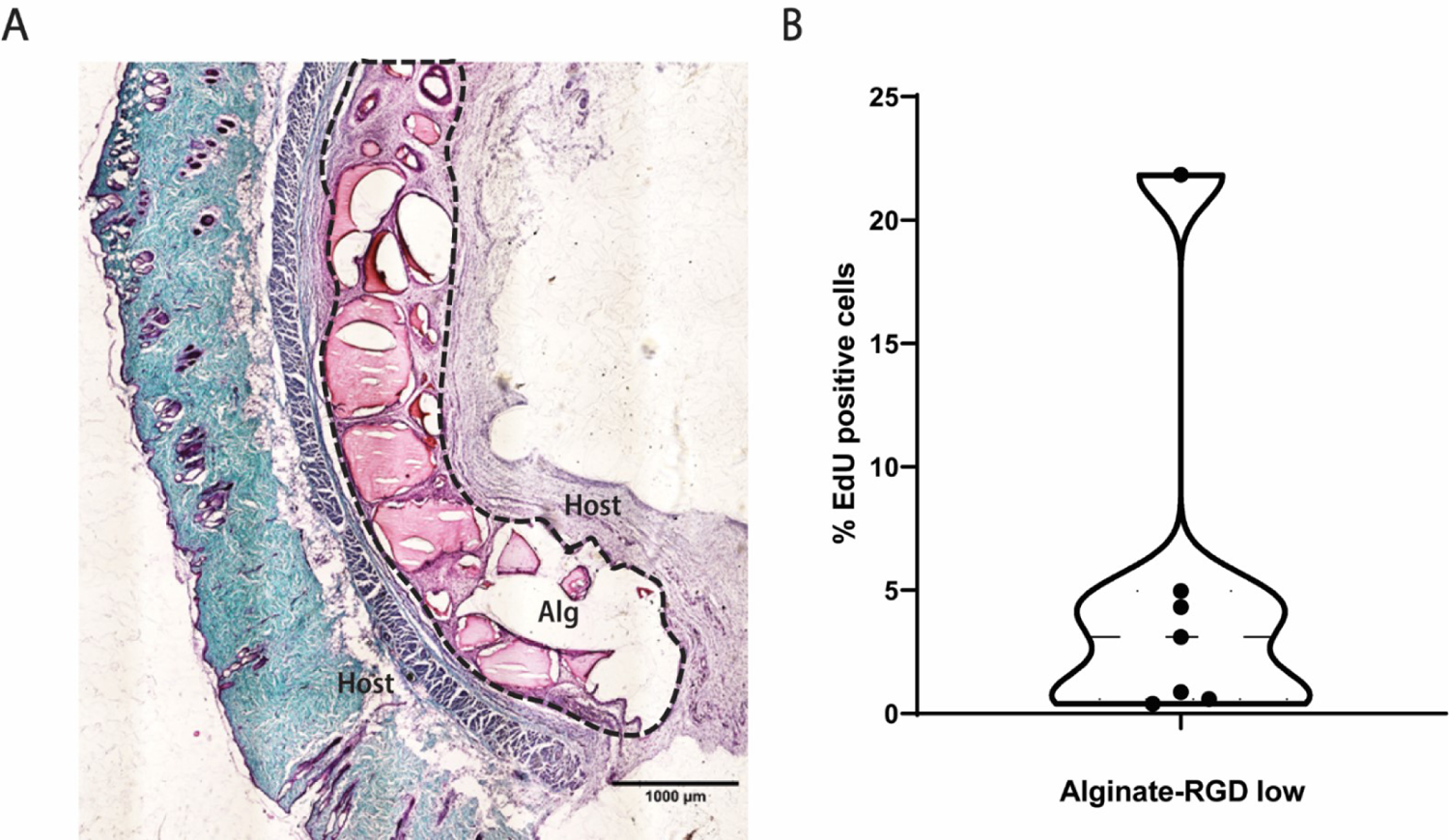
In vivo behavior hMSC encapsulated in alginate-RGD low. (A) Safranin-O staining counterstained with fast green of alginate-RGD low discs implanted in pockets formed on the back of rats for 3 weeks. Alginate-RGD low, (Alg) in red and circled by black dash lines, kept its overall integrity throughout the implantation time. Collagen in green indicates that there was not a thick capsuled formed around the hydrogel. Host stands for the host tissue. n=7. Scale bar=1000 μm (B) Distribution of the percentage of EdU-positive hMSCs inside the implanted alginate-RGD low discs, showing a very low percentage of EdU^+^ cells, indicating that most of the hMSCs encapsulated in alginate-RGD low were quiescent.

**Table S1:**
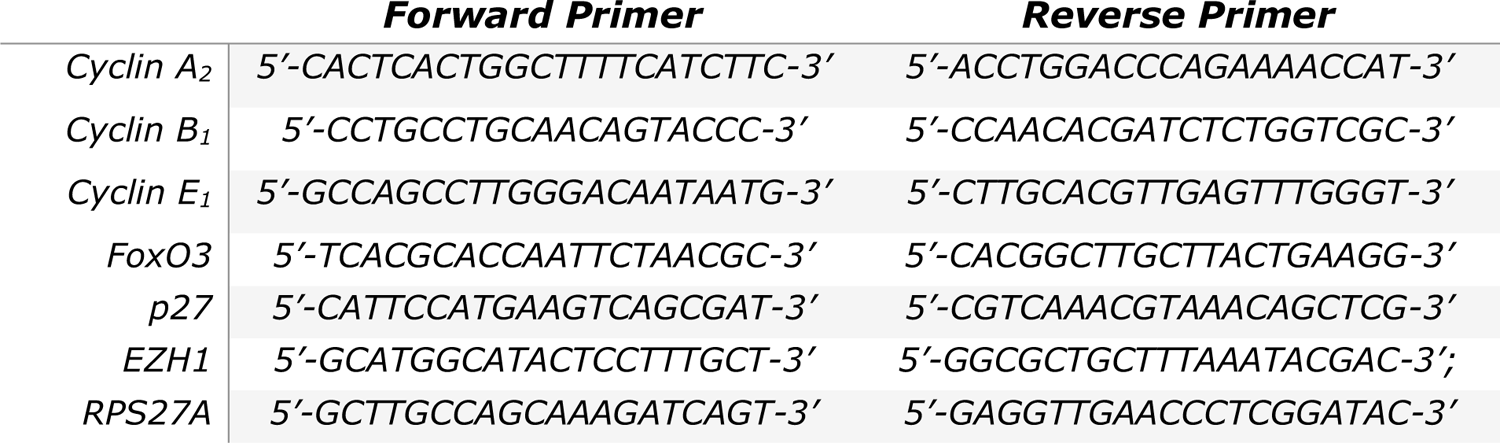
Forward and Reverse primers list

## Notes

### Competing Interest Statement

The authors have declared no competing interest.

## References

1. Biteau, B., C.E. Hochmuth, and H. Jasper, Maintaining tissue homeostasis: dynamic control of somatic stem cell activity. Cell Stem Cell, 2011. 9(5): p. 402–11.

2. Gattazzo, F., A. Urciuolo, and P. Bonaldo, Extracellular matrix: A dynamic microenvironment for stem cell niche. Biochimica Et Biophysica Acta-General Subjects, 2014. 1840(8): p. 2506–2519.

3. Ferreira, M.S., et al., Two-dimensional polymer-based cultures expand cord blood-derived hematopoietic stem cells and support engraftment of NSG mice. Tissue Eng Part C Methods, 2013. 19(1): p. 25–38.

4. Gilbert, P.M., et al., Substrate elasticity regulates skeletal muscle stem cell self-renewal in culture. Science, 2010. 329(5995): p. 1078-81.

5. Yu, N.Y., et al., Live Tissue Imaging to Elucidate Mechanical Modulation of Stem Cell Niche Quiescence. Stem Cells Transl Med, 2017. 6(1): p. 285–292.

6. Winer, J.P., et al., Bone marrow-derived human mesenchymal stem cells become quiescent on soft substrates but remain responsive to chemical or mechanical stimuli. Tissue Eng Part A, 2009. 15(1): p. 147–54.

7. Sun, M.Y., et al., Effects of Matrix Stiffness on the Morphology, Adhesion, Proliferation and Osteogenic Differentiation of Mesenchymal Stem Cells. International Journal of Medical Sciences, 2018. 15(3): p. 257–268.

8. Medeiros Tavares Marques, J.C., et al., Identification of new genes associated to senescent and tumorigenic phenotypes in mesenchymal stem cells. Sci Rep, 2017. 7(1): p. 17837.

9. Yein Lee, Y.A., Jinsung Ahn, Deogil Kim, Seunghee Oh, Donyoung Kang, Hyungsuk Lee, James J. Moon, Bogyu Choi & Soo-Hong Lee, Three-dimensional microenvironmental priming of human mesenchymal stem cells in hydrogels facilitates efficient and rapid retroviral gene transduction via accelerated cell cycle synchronization. NPG Asia Materials, 2019. 11, 27.

10. Bartosh, T.J. and J.H. Ylostalo, Efficacy of 3D Culture Priming is Maintained in Human Mesenchymal Stem Cells after Extensive Expansion of the Cells. Cells, 2019. 8(9).

11. Grayson, W.L., T. Ma, and B. Bunnell, Human mesenchymal stem cells tissue development in 3D PET matrices. Biotechnol Prog, 2004. 20(3): p. 905–12.

12. Huang, G.Y., et al., Engineering three-dimensional cell mechanical microenvironment with hydrogels. Biofabrication, 2012. 4(4).

13. Therese Andersen, P.A.-E., Michael Dornish, 3D Cell Culture in Alginate Hydrogels. Microarrays, 2015. 4(**2**): p. 133–161.

14. Ooi, H.W., et al., Thiol-Ene Alginate Hydrogels as Versatile Bioinks for Bioprinting. Biomacromolecules, 2018. 19(8): p. 3390–3400.

15. Knoll, G.A., et al., Multilayered Short Peptide-Alginate Blends as New Materials for Potential Applications in Cartilage Tissue Regeneration. Journal of Nanoscience and Nanotechnology, 2016. 16(3): p. 2464–2473.

16. Ooi, H.W., et al., Hydrogels that listen to cells: a review of cell-responsive strategies in biomaterial design for tissue regeneration. Materials Horizons, 2017. 4(6): p. 1020–1040.

17. Fonseca, K.B., et al., Injectable MMP-Sensitive Alginate Hydrogels as hMSC Delivery Systems. Biomacromolecules, 2014. 15(1): p. 380–390.

18. Digirolamo, C.M., et al., Propagation and senescence of human marrow stromal cells in culture: a simple colony-forming assay identifies samples with the greatest potential to propagate and differentiate. Br J Haematol, 1999. 107(2): p. 275–81.

19. Neves, S.C., et al., Correction: Biofunctionalized pectin hydrogels as 3D cellular microenvironments. J Mater Chem B, 2015. 3(42): p. 8422.

20. Yao, G., Modelling mammalian cellular quiescence. Interface Focus, 2014. 4(3): p. 20130074.

21. McClatchey, A.I. and A.S. Yap, Contact inhibition (of proliferation) redux. Current Opinion in Cell Biology, 2012. 24(5): p. 685–694.

22. Rumman, M., J. Dhawan, and M. Kassem, Concise Review: Quiescence in Adult Stem Cells: Biological Significance and Relevance to Tissue Regeneration. Stem Cells, 2015. 33(10): p. 2903–2912.

23. Tumpel, S. and K.L. Rudolph, Quiescence: Good and Bad of Stem Cell Aging. Trends in Cell Biology, 2019. 29(8): p. 672–685.

24. Johnson, D.G. and C.L. Walker, Cyclins and cell cycle checkpoints. Annual Review of Pharmacology and Toxicology, 1999. 39: p. 295–312.

25. Cheung, T.H. and T.A. Rando, Molecular regulation of stem cell quiescence. Nature Reviews Molecular Cell Biology, 2013. 14(6): p. 329–340.

26. Sherr, C.J. and J.M. Roberts, Inhibitors of Mammalian G(1) Cyclin-Dependent Kinases. Genes & Development, 1995. 9(10): p. 1149–1163.

27. Renault, V.M., et al., FoxO3 Regulates Neural Stem Cell Homeostasis. Cell Stem Cell, 2009. 5(5): p. 527–539.

28. Hidalgo, I., et al., Ezh1 Is Required for Hematopoietic Stem Cell Maintenance and Prevents Senescence-like Cell Cycle Arrest. Cell Stem Cell, 2012. 11(5): p. 649–662.

29. Campisi, J., Senescent cells, tumor suppression, and organismal aging: Good citizens, bad neighbors. Cell, 2005. 120(4): p. 513–522.

30. Kubota, Y., K. Takubo, and T. Suda, Bone marrow long label-retaining cells reside in the sinusoidal hypoxic niche. Biochem Biophys Res Commun, 2008. 366(2): p. 335–9.

31. Suda, T., K. Takubo, and G.L. Semenza, Metabolic regulation of hematopoietic stem cells in the hypoxic niche. Cell Stem Cell, 2011. 9(4): p. 298–310.

32. Coller, H.A., The paradox of metabolism in quiescent stem cells. FEBS Lett, 2019. 593(20): p. 2817–2839.

33. Read, G.H., et al., Three-dimensional alginate hydrogels for radiobiological and metabolic studies of cancer cells. Colloids Surf B Biointerfaces, 2018. 171: p. 197–204.

34. Kapp, T.G., et al., A Comprehensive Evaluation of the Activity and Selectivity Profile of Ligands for RGD-binding Integrins. Sci Rep, 2017. 7: p. 39805.

35. Barczyk, M., S. Carracedo, and D. Gullberg, Integrins. Cell Tissue Res, 2010. 339(1): p. 269–80.

36. Chen, C.S., et al., Cell shape provides global control of focal adhesion assembly. Biochem Biophys Res Commun, 2003. 307(2): p. 355–61.

37. Ezzell, R.M., et al., Vinculin promotes cell spreading by mechanically coupling integrins to the cytoskeleton. Exp Cell Res, 1997. 231(1): p. 14–26.

38. Gallant, N.D., K.E. Michael, and A.J. Garcia, Cell adhesion strengthening: contributions of adhesive area, integrin binding, and focal adhesion assembly. Mol Biol Cell, 2005. 16(9): p. 4329–40.

39. Jansen, L.E., et al., Mechanics of intact bone marrow. J Mech Behav Biomed Mater, 2015. 50: p. 299–307.

40. Abdallah, B.M. and M. Kassem, Human mesenchymal stem cells: from basic biology to clinical applications. Gene Ther, 2008. 15(2): p. 109–16.

41. Simmons, P.J. and B. Torok-Storb, Identification of stromal cell precursors in human bone marrow by a novel monoclonal antibody, STRO-1. Blood, 1991. 78(1): p. 55-62.

42. Bernal, A. and L. Arranz, Nestin-expressing progenitor cells: function, identity and therapeutic implications. Cell Mol Life Sci, 2018. 75(12): p. 2177–2195.

43. Mendez-Ferrer, S., et al., Mesenchymal and haematopoietic stem cells form a unique bone marrow niche. Nature, 2010. 466(7308): p. 829-34.

44. Xie, L., et al., Characterization of Nestin, a Selective Marker for Bone Marrow Derived Mesenchymal Stem Cells. Stem Cells Int, 2015. 2015: p. 762098.

45. Laplante, M. and D.M. Sabatini, mTOR signaling at a glance. Journal of Cell Science, 2009. 122(20): p. 3589–3594.

46. Sabatini, D.M., Twenty-five years of mTOR: Uncovering the link from nutrients to growth. Proc Natl Acad Sci U S A, 2017. 114(45): p. 11818–11825.

47. Rodgers, J.T., et al., mTORC1 controls the adaptive transition of quiescent stem cells from G0 to G(Alert). Nature, 2014. 510(7505): p. 393-6.

48. Dan, H.C., et al., Akt-dependent Activation of mTORC1 Complex Involves Phosphorylation of mTOR (Mammalian Target of Rapamycin) by I kappa B Kinase alpha (IKK alpha). Journal of Biological Chemistry, 2014. 289(36): p. 25227–25240.

49. Jacinto, E., et al., Mammalian TOR complex 2 controls the actin cytoskeleton and is rapamycin insensitive. Nature Cell Biology, 2004. 6(11): p. 1122–U30.

50. Breuleux, M., et al., Increased AKT S473 phosphorylation after mTORC1 inhibition is rictor dependent and does not predict tumor cell response to PI3K/mTOR inhibition. Molecular Cancer Therapeutics, 2009. 8(4): p. 742–753.

51. Vincent, E.E., et al., Akt phosphorylation on Thr308 but not on Ser473 correlates with Akt protein kinase activity in human non-small cell lung cancer. British Journal of Cancer, 2011. 104(11): p. 1755–1761.

52. Vadlakonda, L., M. Pasupuleti, and R. Pallu, Role of PI3K-AKT-mTOR and Wnt Signaling Pathways in Transition of G1-S Phase of Cell Cycle in Cancer Cells. Front Oncol, 2013. 3: p. 85.

53. Morikawa, S., et al., Prospective identification, isolation, and systemic transplantation of multipotent mesenchymal stem cells in murine bone marrow. J Exp Med, 2009. 206(11): p. 2483–96.

54. Sage, J., The retinoblastoma tumor suppressor and stem cell biology. Genes Dev, 2012. 26(13): p. 1409–20.

55. Cho, I.J., et al., Mechanisms, Hallmarks, and Implications of Stem Cell Quiescence. Stem Cell Reports, 2019. 12(6): p. 1190–1200.

56. Katoh, K., Y. Kano, and Y. Noda, Rho-associated kinase-dependent contraction of stress fibres and the organization of focal adhesions. J R Soc Interface, 2011. 8(56): p. 305–11.

57. Olson, E.N. and A. Nordheim, Linking actin dynamics and gene transcription to drive cellular motile functions. Nat Rev Mol Cell Biol, 2010. 11(5): p. 353–65.

58. Kovacs, M., et al., Mechanism of blebbistatin inhibition of myosin II. J Biol Chem, 2004. 279(34): p. 35557–63.

59. Boyette, L.B. and R.S. Tuan, Adult Stem Cells and Diseases of Aging. J Clin Med, 2014. 3(1): p. 88–134.

60. Prockop, D.J., I. Sekiya, and D.C. Colter, Isolation and characterization of rapidly self-renewing stem cells from cultures of human marrow stromal cells. Cytotherapy, 2001. 3(5): p. 393–396.

61. Moya, A., et al., Quiescence Preconditioned Human Multipotent Stromal Cells Adopt a Metabolic Profile Favorable for Enhanced Survival under Ischemia. Stem Cells, 2017. 35(1): p. 181–196.

62. Chen, M., et al., Serum starvation induced cell cycle synchronization facilitates human somatic cells reprogramming. PLoS One, 2012. 7(4): p. e28203.

63. Morrison, S.J. and D.T. Scadden, The bone marrow niche for haematopoietic stem cells. Nature, 2014. 505(7483): p. 327-334.

64. Ehninger, A. and A. Trumpp, The bone marrow stem cell niche grows up: mesenchymal stem cells and macrophages move in. Journal of Experimental Medicine, 2011. 208(3): p. 421–428.

65. Xu, J., et al., Chondrogenic differentiation of human mesenchymal stem cells in three-dimensional alginate gels. Tissue Eng Part A, 2008. 14(5): p. 667–80.

66. Maia, F.R., et al., Effect of cell density on mesenchymal stem cells aggregation in RGD-alginate 3D matrices under osteoinductive conditions. Macromol Biosci, 2014. 14(6): p. 759–71.

67. Ma, H.L., et al., Chondrogenesis of human mesenchymal stem cells encapsulated in alginate beads. J Biomed Mater Res A, 2003. 64(2): p. 273–81.

68. Cheung, T.H. and T.A. Rando, Molecular regulation of stem cell quiescence. Nat Rev Mol Cell Biol, 2013. 14(6): p. 329–40.

69. Miyamoto, K., et al., Foxo3a is essential for maintenance of the hematopoietic stem cell pool. Cell Stem Cell, 2007. 1(1): p. 101–112.

70. Gopinath, S.D., et al., FOXO3 Promotes Quiescence in Adult Muscle Stem Cells during the Process of Self-Renewal. Stem Cell Reports, 2014. 2(4): p. 414–426.

71. Zou, P., et al., p57(KiP2) and p27(Kip1) Cooperate to Maintain Hematopoietic Stem Cell Quiescence through Interactions with Hsc70. Cell Stem Cell, 2011. 9(3): p. 247–261.

72. Oki, T., et al., A novel cell-cycle-indicator, mVenus-p27K(-), identifies quiescent cells and visualizes G0-G1 transition. Scientific Reports, 2014. 4.

73. Rumman, M., et al., Induction of quiescence (G0) in bone marrow stromal stem cells enhances their stem cell characteristics. Stem Cell Res, 2018. 30: p. 69–80.

74. Fujimaki, K., et al., Graded regulation of cellular quiescence depth between proliferation and senescence by a lysosomal dimmer switch. Proceedings of the National Academy of Sciences of the United States of America, 2019. 116(45): p. 22624–22634.

75. Chiu, H.Y., Y.G. Tsay, and S.C. Hung, Involvement of mTOR-autophagy in the selection of primitive mesenchymal stem cells in chitosan film 3-dimensional culture. Sci Rep, 2017. 7(1): p. 10113.

76. Barrientos, S., et al., Growth factors and cytokines in wound healing. Wound Repair Regen, 2008. 16(5): p. 585–601.

77. Ferreira, S.A., et al., Neighboring cells override 3D hydrogel matrix cues to drive human MSC quiescence. Biomaterials, 2018. 176: p. 13–23.

